# Structurally distinct endocytic pathways for B cell receptors in B lymphocytes

**DOI:** 10.1101/394015

**Authors:** Aleah D. Roberts, Thaddeus M. Davenport, Andrea M. Dickey, Regina Ahn, Kem A. Sochacki, Justin W. Taraska

## Abstract

B lymphocytes play a critical role in adaptive immunity. Upon antigen binding, B cell receptors (BCR) cluster on the plasma membrane and are internalized by endocytosis. In this process, B cells capture diverse antigens in various contexts and concentrations. However, it is unclear whether the mechanism of BCR endocytosis changes in response to these factors. Here, we studied the mechanism of soluble antigen-induced BCR clustering and internalization in a cultured human B cell line using correlative super resolution fluorescence and platinum replica electron microscopy. First, by visualizing nanoscale BCR clusters, we provide direct evidence that BCR cluster size increases with F(ab’)2 concentration. Next, we show that the physical mechanism of internalization switches in response to BCR cluster size. At low concentrations of antigen, B cells internalize small BCR clusters by classical clathrin-mediated endocytosis. At high antigen concentrations, when clusters size increases beyond the size of a single clathrin coated pit, B cells retrieve receptor clusters using large invaginations of the plasma membrane capped with clathrin. At these sites, we observed early and sustained recruitment of actin and an actin polymerizing protein FCHSD2. We further show that actin recruitment is required for the efficient generation of these novel endocytic carriers and for their capture into the cytosol. We propose that in B cells, the mechanism of endocytosis switches to accommodate large receptor clusters formed when cells encounter high concentrations of soluble antigen. This mechanism is regulated by the organization and dynamics of the cortical actin cytoskeleton.

## INTRODUCTION

B lymphocytes are critical in adaptive immune responses to pathogens (Trombetta and Mellman, 2005). Dysregulation of B cell activity can lead to cancer, autoimmunity, and allergy (Avalos et al., 2014). The B cell receptor (BCR), abundant in the plasma membrane of B cells, binds to and internalizes antigens by endocytosis for processing and presentation to T cells (Clark et al., 2004; Harwood and Batista, 2010). BCR activation, along with T cell engagement, promotes the expansion of a clonal lineage of antigen-specific B cells. As a first step, clustering of BCRs in response to antigen binding initiates a signaling cascade that triggers the internalization of receptor/antigen complexes by endocytosis (Harwood and Batista, 2010; Pierce and Liu, 2010). Antigen structure, concentration, and presentation may all influence the size of BCR/antigen clusters formed on the plasma membrane (Batista et al., 2001; Srinivas Reddy et al., 2011; Thyagarajan et al., 2003). The radii of these clusters can vary from ~60 nm to over 1 micron (Lee et al., 2017; Pierce and Liu, 2010; Stone et al., 2017). The membrane trafficking pathways that enable B cells to specifically internalize antigen in spite of such structural variability are unclear.

Clathrin-mediated endocytosis (CME) is the primary mechanism of endocytosis in eukaryotic cells (Conner and Schmid, 2003; McMahon and Boucrot, 2011; Mettlen et al., 2018). In CME, a complex of adaptors, accessory proteins, lipids, and cargo recruit clathrin to the inner leaflet of the plasma membrane where it assembles as a lattice that curves and drives the formation of vesicles (McMahon and Boucrot, 2011; Mettlen et al., 2018). Clathrin has been shown to play an important role in BCR endocytosis and BCR signaling (Natkanski et al., 2013; Salisbury et al., 1980; Stoddart et al., 2005). For example, knockdown of clathrin dramatically impairs BCR internalization (Stoddart et al., 2005). Similarly, mutations of clathrin-adaptor domains found in the BCR complex results in active receptors that accumulate at the plasma membrane and are observed in lymphomas (Davis et al., 2010). The precise role of clathrin in BCR endocytosis, however, is not well defined.

While clathrin has been shown to be important for BCR endocytosis, B cells can retrieve BCRs in the absence of clathrin (Stoddart et al., 2005). This raises the possibility that there may be other mechanisms of BCR internalization. A number of alternative endocytic pathways have been described in eukaryotic cells (Mayor et al., 2014; Watanabe and Boucrot, 2017). These clathrin-independent endocytosis (CIE) pathways include micron-scale pinocytosis and phagocytosis processes as well as nanometer-scale mechanisms such as caveolae, flotillin-based, endophilin-based, actin-based, and other less well-defined vesicle or transport processes (Ferreira and Boucrot, 2018; Mayor et al., 2014). Thus, it is not known if CIE uptake pathways exist for B cell receptors or if a unique hybrid form of endocytosis operates during antigen-mediated BCR internalization. In support of other pathways, previous work has shown that B cells are capable of internalizing large cargos through a process akin to phagocytosis (Souwer et al., 2009). This capacity for phagocytic-like processes is consistent with observations that actin and myosin play an important role in antigen-mediated BCR clustering, activation, and internalization (Brown and Song, 2001; Tolar, 2017). Furthermore, large pieces of B cell and target membranes are internalized when B cells interact with antigen presented on supported membranes (Natkanski et al., 2013). Interestingly, this membrane capture process depends on clathrin, but doesn’t appear to wholly resemble classic clathrin-mediated endocytosis in size, structure, or molecular composition (Natkanski et al., 2013).

How antigen concentration and receptor cluster size impacts the proposed pathways of BCR endocytosis are also unclear. Concentration-dependent internalization mechanisms have been proposed for some receptors such as epidermal growth factor receptor (EGFR)(Tomas et al., 2014). Low concentrations of epidermal growth factor (EGF) stimulate EGFR uptake by a clathrin-dependent mechanism, but at high EGF concentrations internalization occurs by a mixture of clathrin-dependent and clathrin–independent mechanisms (Sigismund et al., 2008). Such behaviors have not been described for the BCR. However, B cells might benefit from such a concentration-dependent switch in the mechanism of BCR endocytosis. Specifically, by adjusting the endocytic mechanism of BCR uptake in response to antigen concentrations, B cells might be better able to rapidly and appropriately respond to a wide-range of changes in the concentration of foreign antigens during an infection. Likewise, B cells could avoid inappropriate or unnecessary responses to common self-antigens (e.g. leading to autoimmunity) or transient exposures to foreign but non-pathogenic molecules (e.g. leading to allergy). However, no systematic study has been made to directly visualize the nanoscale structural changes in the plasma membrane of B cells during antigen challenge.

Here, we study the formation and structure of anti-human IgM F(ab’)2-induced endocytic carriers in the IgM+ DG-75 human B cell line to understand the mechanism of BCR internalization. Specifically, we investigated the role of clathrin in mediating BCR endocytosis across a range of antigen concentrations. First, we measure BCR internalization in live cells and observe the nanoscale structure and dynamics of antigen-induced BCR clustering and internalization in real time. Next, we use correlative super resolution fluorescence and platinum replica transmission electron microscopy (CLEM) to directly map the nanoscale architecture of BCR endocytic structures throughout the process of antigen stimulation and uptake. We show that BCR clusters induced by increasing concentrations of antigen and time are captured by distinct nanoscale plasma membrane invaginations capped with clathrin. We then show that these distinct plasma membrane structures recruit actin and that actin polymerization is important for the capture and internalization of large BCR clusters. This work provides direct evidence for the structural complexity of BCR endocytosis and offers direct insights into a novel endocytic mechanism that allows B cells to tailor their endocytic machinery to accommodate the wide range of BCR cluster sizes formed at the plasma membrane of living B cells.

## RESULTS

### Antigen stimulation across a wide range of concentrations induces efficient BCR internalization

First, to characterize the temporal and spatial dynamics of BCR internalization in our system, we imaged live DG-75 cells with confocal microscopy (Figure 1a-b). Cross-linking anti-human IgM F(ab’)2 antibody was added to DG-75 cells to a final concentration of 12 μg/mL at 37°C to induce activation and internalization of native BCRs pre-labeled at a low concentration with fluorescent single chain F(ab) fragments. Five minutes after the addition of the crosslinking ligand, cell-surface BCRs formed small fluorescent patches that condensed into larger clusters that were internalized into the cytoplasm over a period of 25 minutes (Figure 1a-b). These results are consistent with previous studies of BCR clustering (Brown and Song, 2001). Furthermore, population-based fluorescence assisted cell sorting (FACS) measurements of receptor internalization indicated that BCRs were internalized across a wide range of stimulating antigen concentration (0.88 μg/ml-14 μg/ml F(ab’)2) (Supplementary figure 1 a, b and c). Here, similar amounts of fluorescently-labeled BCR-bound F(ab) fragments were internalized at both low (2 μg/ml F(ab’)2) and high (8 μg/ml F(ab’)2) concentrations of simulating antigen (Supplementary figure 2 d). These results were supported by thin section transmission electron microscope (TEM) of B cells stimulated in the presence of membrane-bound ferritin, a small electron-dense indicator that labels the outer leaflet of the plasma membrane and inner leaflet of endocytic vesicles. In unstimulated cells little internalization of surface-bound ferritin was observed by thin section TEM (Supplementary figure 2 a-c), but ferritin-positive endocytic structures were observed attached to the plasma membrane and deeper in the cytosol after cells were stimulated for 15 minutes with 8 μg/mL anti-human IgM F(ab’)2 (Supplementary figure 2 d and e).

**Figure 1.**
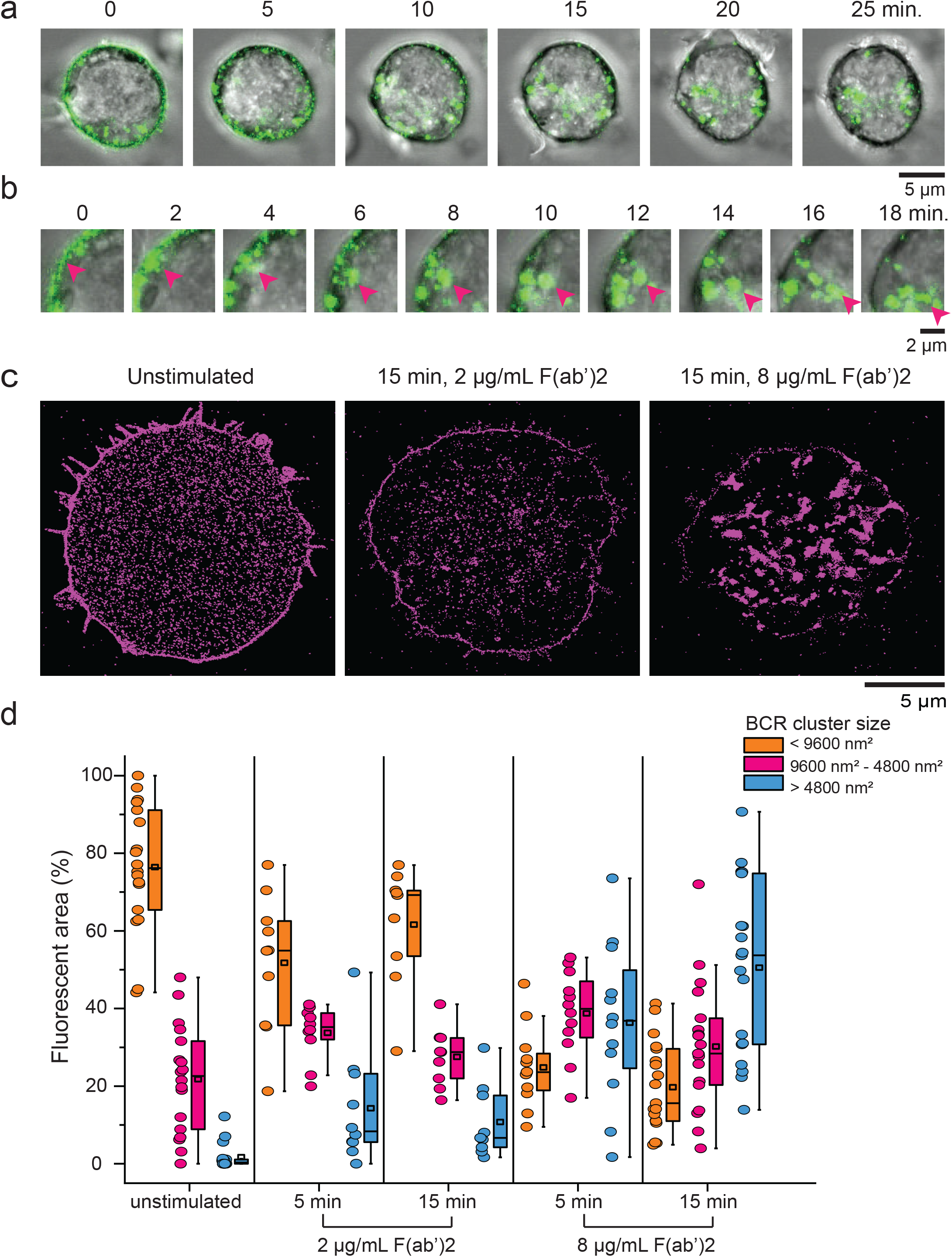
F(ab’)2 stimulation of DG-75 cells induces BCR clustering and internalization. (a) Confocal images of a live DG-75 cell labeled with anti-human IgM F(ab)-Alexa Fluor 488 and stimulated with 12 μg/ml of anti-human IgM F(ab’)2 at t = 0 min. BCR is shown in green and transmitted light is grayscale. (b) Upper left region of cell in (a); pink arrow tracks BCRs that cluster and internalize into the cytoplasm. (c) Reconstructed super-resolution images of the BCR (magenta) clustering on the bottom surface of DG-75 cells labeled with anti-human IgM F(ab)-Alexa Fluor 647 and left unstimulated or stimulated with anti-human IgM F(ab’)2 at either 2 or 8 μg/mL F(ab’)2 for 15 min prior to plating, fixation, and imaging. (d) Percent of total fluorescent area in each cell composed of small (<9600 nm2, orange bar), intermediate (9600-48000 nm2, pink bar), and large (> 48000 nm2, blue bar) punctae for unstimulated DG-75 cells, cells stimulated with 2 μg/mL F(ab’)2 for 5 or 15 minutes, and cells stimulated with 8 μg/mL F(ab’)2 for 5 or 15 minutes (n = unstimulated 18 cells – 47,847 spots, 5 min 2 μg/ml 10 cells – 23,728 spots, 5 min 8 μg/ml 12 cells – 10,497 spots, 15 min 2 μg/ml 9 cells – 14,347 spots, 15 min 8 μg/ml 19 cells – 8,237 spots). Plots show the mean (square), median (line), 25/75 percentile range (box), and outliers with a coefficient value of 1.5 and data points (circles) from each cell.

### BCR cluster size is dependent on concentration of F(ab’)2 stimulation

As a first step in BCR uptake, antigen-bound BCRs aggregate on the plasma membrane (Lee et al., 2017). To determine whether the concentration of soluble ligand alters the nanoscale distribution and clustering of BCRs in the plasma membrane, we measured the size of BCR clusters using direct stochastic optical reconstruction microscopy (dSTORM), a super-resolution localization microcopy method (Betzig et al., 2006; Heilemann et al., 2008). Figure 1c shows representative dSTORM images of DG-75 cells stained with Alexa 647-labeled anti-human IgM F(ab) fragments under three conditions: unstimulated, incubated with low (2 μg/mL), or high (8 μg/mL) concentrations of F(ab’)2 for 15 minutes at 37 °C before attachment to coverslips and fixation. In unstimulated cells, BCR fluorescence was uniformly distributed across the plasma membrane with few measurable clusters (Figure 1c left). After 15 minutes of stimulation with 2 μg/mL F(ab’)2, the size of BCR fluorescent spots increased (Figure 1c center). However, the most dramatic change in BCR distribution was observed after 15 minutes of stimulation with 8 μg/mL F(ab’)2, which resulted in very large clusters of BCR fluorescence (Figure 1c right).

As shown in figure 1, after BCRs cluster they are endocytosed into the cytosol. To investigate the mechanism of endocytosis, we first classified these BCR clusters into three groups based on their size relative to one candidate endocytic carrier, clathrin coated pits. Clathrin coated pits have an area of around 18,000 nm^2^ on average, but range from 10,000 nm^2^ to 50,000 nm^2^ in area, or 55 to 125 nm in radius (Dambournet et al., 2018; Sochacki et al., 2017; Sochacki et al., 2012). Thus, using these sizes as a reference, we characterized: 1) small spots (<9600 nm^2^), which are smaller than a typical clathrin coated pit; 2) intermediate spots (9600-48000 nm^2^), which are comparable in size to the majority of clathrin coated pits; and 3) large fluorescent spots (>48000 nm^2^), which are over 2.5 times the mean area of clathrin coated pits. Approximately 76% of the fluorescent area in unstimulated cells was composed of small spots, with a majority of the remaining made up of spots between 9600-48000 nm^2^ (Figure 1d). Large clusters made up an insignificant portion (around 2%) of the fluorescence in unstimulated cells. In contrast, for cells treated with 8 μg/mL F(ab’)2, large fluorescent spots made up an increasing large proportion of the fluorescent area (36% and 51%) at both the 5 and 15 minute time points respectively (Figure 1d). This was not the case for cells treated with 2 μg/mL F(ab’)2, in which 62% of the fluorescent area was made up of small spots. Although there was a slightly increased proportion of intermediate and large fluorescent clusters relative to unstimulated cells (Figure 1d). Thus, increasing the concentration of soluble F(ab’)2 dramatically increased the size of BCR clusters on the plasma membrane. Of note, after 15 minutes of stimulation with 8 μg/mL F(ab’)2, we predominantly observed very large BCR clusters that exceeded the size of a typical clathrin coated pit. If BCR clusters exceed the size of their proposed endocytic clathrin carriers, we next asked how these large clusters are captured and internalized.

### Large BCR clusters are associated with distinct endocytic structures at the plasma membrane

Correlative light and electron microscopy (CLEM) provides a unique nanoscale view of the cellular environment of proteins localized by fluorescence microscopy (Kopek et al., 2017; Sochacki et al., 2014). Figure 2a shows correlated super-resolution fluorescence and platinum replica EM image (CLEM image) of the inner plasma membrane of an unstimulated DG-75 B cell plasma membrane. Similar to the data shown in Figure 1c, BCR fluorescence in unstimulated cells was characterized by localizations distributed randomly across the plasma membrane. In platinum replica EM, the structure of plasma membrane-associated actin and vesicles are visible. With higher magnification, no obvious association of the receptor with cytoskeletal filaments or vesicles was seen (Figure 2b). Of note, cells generally exhibited a macroscopic ring pattern of membrane-associated actin. The CLEM images in Figure 2c-f show the inner membrane of DG-75 cells after 5 minutes of stimulation with 2 or 8 μg/mL F(ab’)2. Unlike cells treated with 2 μg/mL F(ab’)2 (Figure 2c and d), after 5 minutes of stimulation with 8 μg/mL F(ab’)2, BCR clusters began to form, and these clusters were often associated with honeycomb clathrin lattices in the correlated TEM image (Figure 2e and f; yellow arrows point to clathrin lattices with BCR fluorescence). After 15 minutes of stimulation with 8 μg/mL F(ab’)2, BCR clusters were larger and were frequently correlated with dense honeycomb arrays that were often associated with distinctive smooth, raised regions of the plasma membrane (Figure 2i and j; yellow arrow points to distinctive smooth raised structure with clathrin and BCR fluorescence). In contrast, cells treated with 2 μg/mL F(ab’)2 showed much smaller BCR fluorescence punctate and few morphological changes of the plasma membrane after 15 minutes (Figure 2g and h).

**Figure 2.**
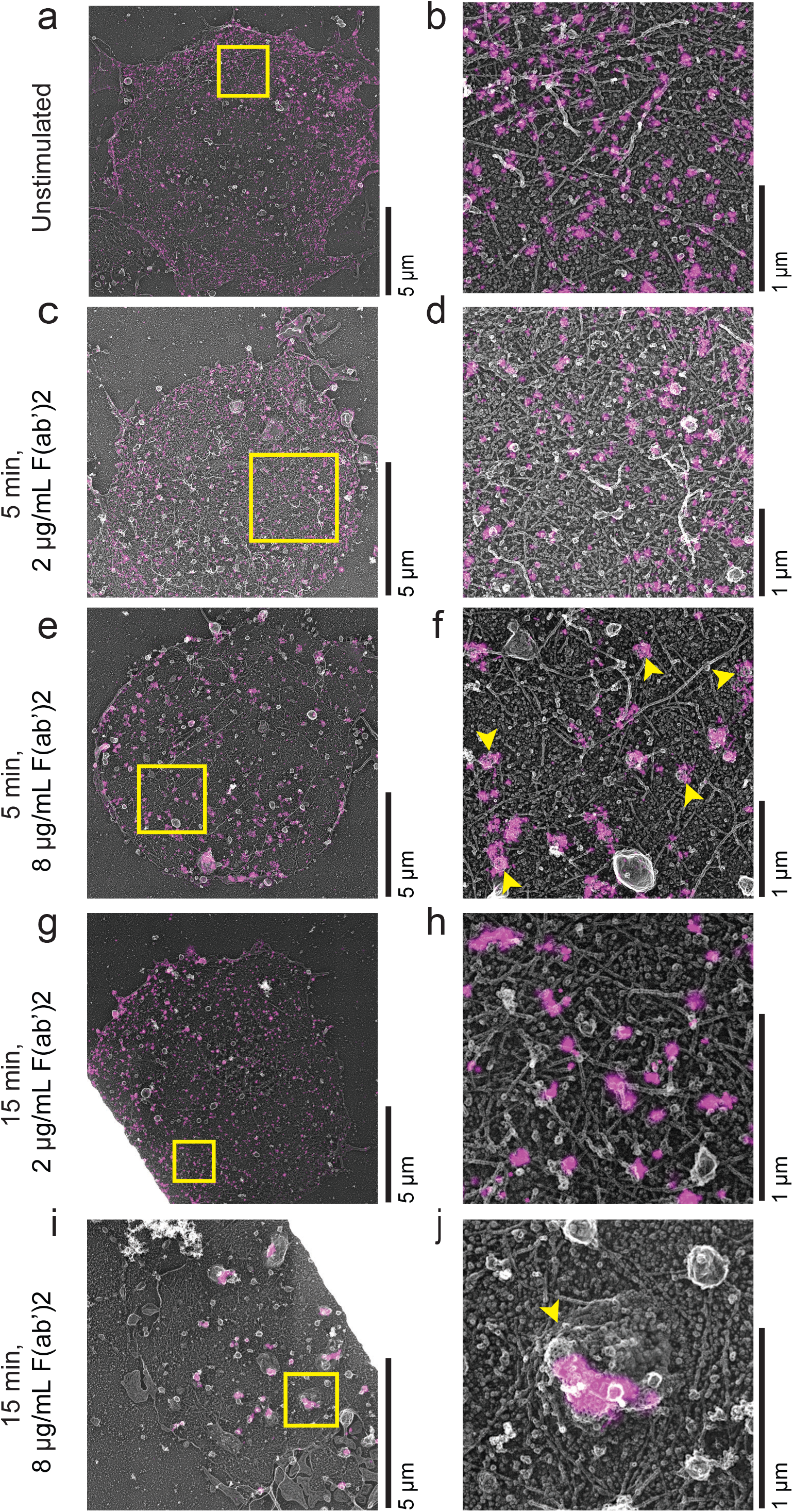
Correlative light and electron microscopy (CLEM) analysis of membrane changes during BCR clustering in DG-75 cells after stimulation. CLEM overlay images of DG-75 cells that have been unroofed and fixed before dSTORM and TEM imaging. For dSTORM imaging cells were labeled with anti-human IgM F(ab)-Alexa Fluor 647. After d-STORM imaging cells were prepared by PREM and imaged using a TEM microscope. Images in parts a, c, e, g and i show the whole cell, and corresponding zoomed in images of the regions in yellow boxes are shown in parts b, d, f, h, and j. The yellow arrows in part f point out regions of the plasma membrane that have BCR fluorescence associated with clathrin lattices. The yellow arrow in part j points out a smooth raised portion of the plasma membrane that has clathrin lattices and BCR fluorescence on it.

To quantitatively track the changes in the membrane and organelles following antigen stimulation in B cells, we first analyzed the structure of regions of the plasma membrane in platinum replica TEM images (Figure 3). For this analysis we focused on identifying distinctive plasma membrane structures in stimulated cells. TEM images were segmented for two major objects: 1) honey-comb clathrin lattices (“Clathrin”) (Figure 3a) and 2) smooth, raised sections of the plasma membrane (“Smooth Raised Membrane”, SRM) (Figure 3b). These two structures often colocalized as SRMs coated with clathrin (clathrin on SRMs) (Figure 3c). Figure 3d-f show an example platinum replica TEM image (Figure 3d) and its corresponding segmented masks identifying clathrin (Figure 3e), SRM (Figure 3f), and clathrin on SRM structures (Figure 3g). Segmented masks were generated for each cell and used to quantitatively analyze the density of each structure on the plasma membrane (Figure 3h-j).This analysis shows that the density of clathrin-coated structures (CCS) is very low in unstimulated DG-75 cells (0.31 CCS/μm^2^ +/− 0.05 SEM) relative to other mammalian cell types (Figure 3h). For example, PC12 cells have ~1.6 CCS/μm^2^ and Hela cells have ~0.7 CCS/μm^2^ (Sochacki et al., 2012). Treatment with 8 μg/mL F(ab’)2 dramatically increased the amount of clathrin on the plasma membrane: at the 5 minute time point we measured 1.32 CCS/μm^2^ +/− 0.2 SEM, and 1.73 CCS/μm2 +/− 0.19 SEM after 15 minutes. There was less recruitment of clathrin to the plasma membrane following treatment with 2 μg/mL F(ab’)2: at 5 minutes we measured a density of 0.65 CCS/μm^2^ +/− 0.11 SEM that only mildly increased at 15 minutes (0.79 CCS/μm^2^ +/− 0.16 SEM) (Figure 3h). Of note, individual clathrin structures did not change in size, with the average Feret size of clathrin structures remaining constant across stimulations (Supplemental Figure 3a). But individual spots did group together, showing a closer nearest-neighbor distance than would be expected from the increased density of clathrin observed over the course of activation (Supplemental Figure 3b). Figure 3i shows that the density of SRMs also increased during stimulation with the largest increase in SRMs seen after 15 minutes in cells exposed to 8 μg/mL of F(ab’)2. Like clathrin, these structures did not change in overall diameter (Supplemental Figure 3c) and yet different than clathrin, showed only moderate clustering (Supplemental Figure 3d). Figure 3j shows that in many cases, clathrin coated structures directly associated with SRMs. The most dramatic overlap of clathrin and SRMs was observed in cells treated for 15 minutes with 8 μg/mL F(ab’)2. Thus, high concentrations of F(ab’)2 induce the formation of large smooth invaginations of the plasma membrane that contain clusters of small clathrin-coated structures.

**Figure 3.**
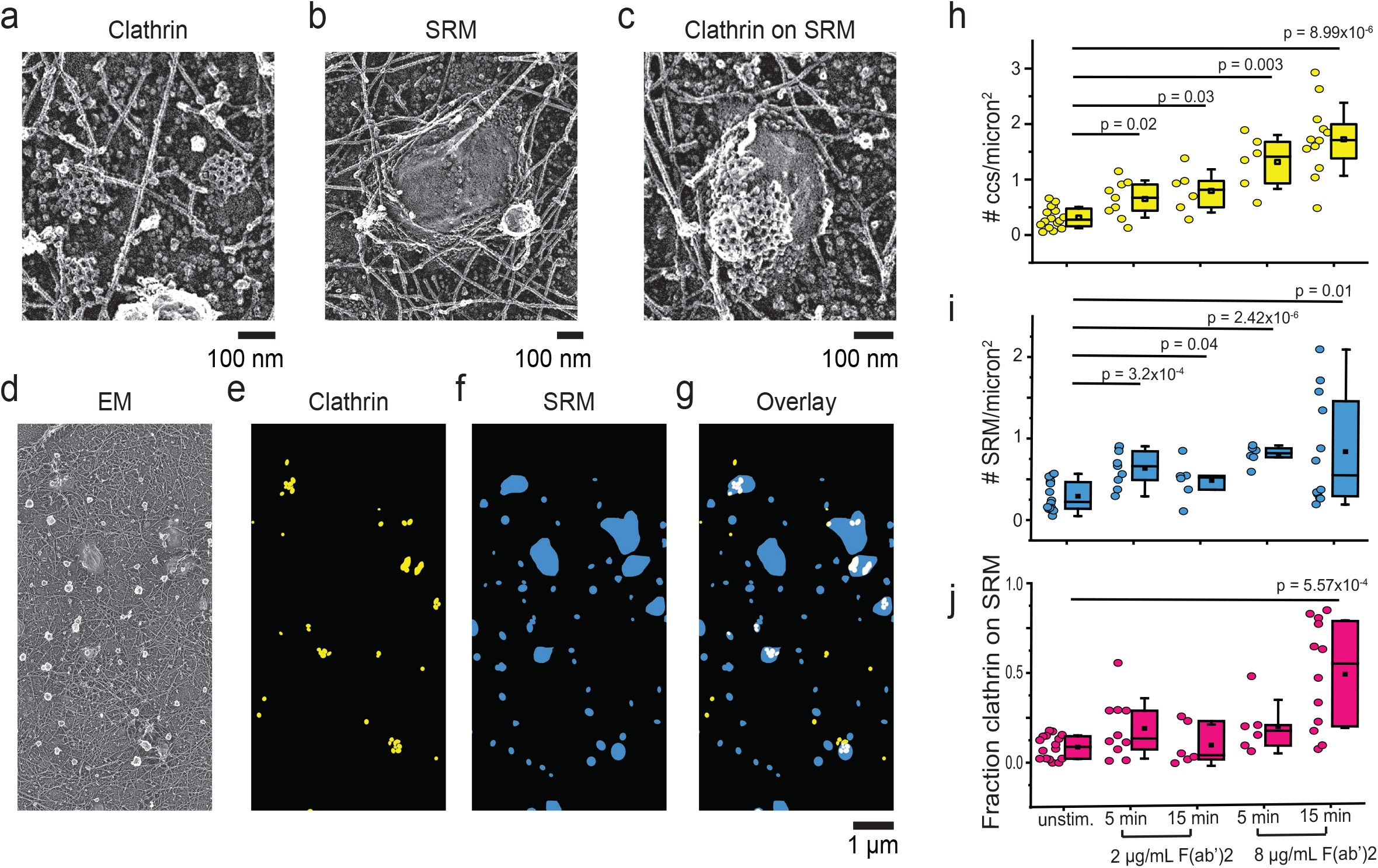
Changes in clathrin and smooth raised membrane structures in TEM images following F(ab’)2 stimulation. Representative PREM images of clathrin (a), smooth raised membrane (SRM) structures (b), and clathrin on SRMs (c). (d) Example PREM image used for identification and resulting segmentation shown for (e) clathrin marked in yellow, (f) smooth raised membrane structures as blue objects, and (g) an overlay with the clathrin found on SRM structures as white regions. These segmented masks were used to measurethe change in the density of CCS (h) and SRM (i) on the membrane. Association of CCS and SRM structures were quantified in (j). Box and whisker plots (h-j) show the mean (square), median (line), 25/75 percentile range (box), and outliers with a coefficient value of 1.5.

### BCR clusters are located on distinct clathrin on SRM structures

The above analysis was performed on platinum replica TEM images to understand the impact of antigen activation on the morphology and structure of the plasma membrane. Next, we quantified changes in the distribution of BCR fluorescence on the plasma membrane relative to these EM-identified locations using super-resolution CLEM images. As seen in the CLEM images (Figure 4a-c), BCR fluorescence did not always exclusively overlap with clathrin but was often found both on clathrin and in the areas closely surrounding clathrin. Thus, using the segmented objects shown in Figure 4 to identify both the directly bound and closely associated fluorescence, we measured the fraction of BCR fluorescence within a 200 nm radius of three different segmented plasma membrane regions (clathrin, SRM, and clathrin located on SRMs) in 5 different experimental conditions (Unstimulated, 5 min 2 μg/mL F(ab’)2, 5 min 8 μg/mL F(ab’)2, 15 min 2 μg/mL F(ab’)2, and 15 min 8 μg/mL F(ab’)2). Figure 4d-f shows that in unstimulated cells, few BCR fluorescent localizations were associated with clathrin, SRMs, or clathrin on SRMs. However, stimulation with 8 μg/mL F(ab’)2 for 5 minutes resulted in a concentration of BCR fluorescence within 200 nm of both clathrin and SRM structures, which further increased at 15 minutes. As shown in Figure 3, under these conditions, most clathrin is found grouped on SRMs, and thus, our measurements in Figure 4 f likewise show that the majority of BCR fluorescence is concentrated on and around these SRM-localized clathrin clusters. In contrast, cells treated with 2 μg/mL F(ab’)2 showed less association of BCR fluorescence with SRM-associated clathrin (Figure 4f). It should be noted that there was little clathrin observed on SRMs under these conditions. Another point of interest in these data is that when cells are stimulated at 8 μg/mL F(ab’)2, clathrin is more highly localized near BCR clusters at the 5 minute time point than SRM structures. SRM structures are not similarly associated with BCR clusters until between 5 and 15 minutes after stimulation. This suggests that clathrin is recruited to BCR clusters before smooth raised membrane structures are generated.

**Figure 4.**
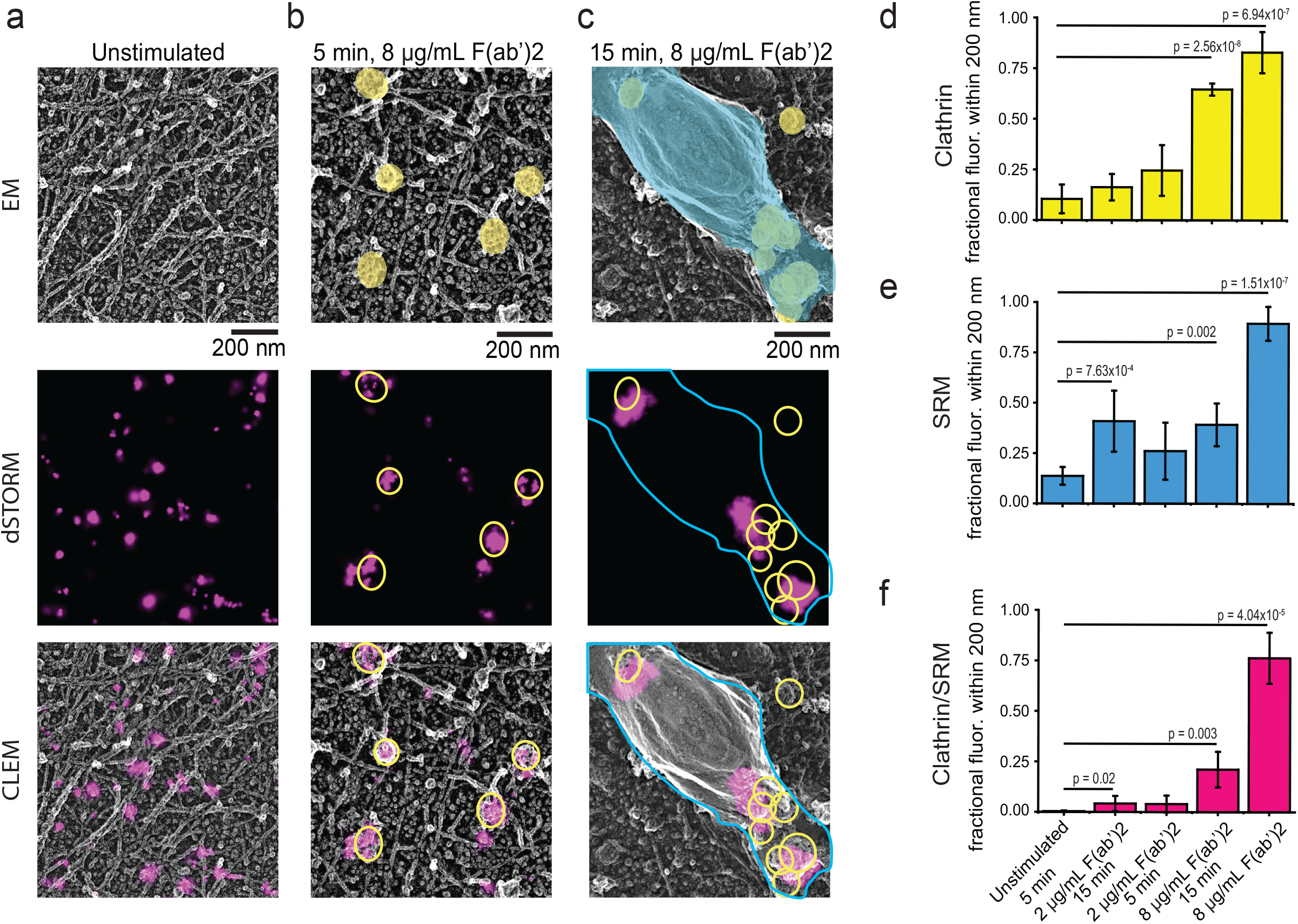
Analysis of changes in the distribution of BCR fluorescence relative to CCS and SRM structures in CLEM images following F(ab’)2 stimulation. Example CLEM images (gray, EM; magenta, dSTORM; and overlay) of (a) unstimulated, (b) 5 min sample, and (c) 15 minutes sample of cells stimulated with 8 μg/mL F(ab’)2. Plots of the fraction of BCR fluorescence measured in an 1800 nm total radius located within 200 nm of a (d) CCS (e) SRM or (f) Clathrin on SRM for 5 different experimental conditions measured from CLEM images. After 5 minutes of stimulation with 8 μg/mL F(ab’)2, approximately 70% of all BCR fluorescence is found within 200 nm of CCS, and after 15 minutes of stimulation with 8 μg/mL of F(ab’)2 nearly 80% of all BCR fluorescence is found within 200 nm of clathrin on SRM structures. Bar graph shows mean +/− SD (n = Unstim. 5 cells, 5 min 2 μg/ml 9 cells, 5 min 8 μg/ml 6 cells, 15 min 2 μg/ml 6 cells, 15 min 8 μg/ml 6 cells).

### Actin is recruited early to BCR clusters stimulated with a high concentration of F(ab’)2

To identify proteins involved in generation of the novel clathrin on SRM structures we used TIRF microscopy to study the colocalization of a panel of candidate endocytic proteins (clathrin, actin, endophilins A1 and A2, synaptojanin 2, and FCHSD2) with BCRs when cells were stimulated with 2 or 8 μg/mL F(ab’)2 (Supplemental Figure 4). Based on our previous data, at the lower concentration of stimulating antigen, BCRs are localized in single clathrin coated structures, and at the high concentrations of antigen, BCR clusters are predominantly localized in and around clustered clathrin on SRM structures. Our data show that both actin and Endophilin A1 were significantly recruited to BCR clusters stimulated with 8 μg/mL F(ab)’2 at the 15 minute time point. Endophilin A1 is thought to be expressed predominantly in neurons(Ringstad et al., 1997). We, however, could not detect endogenous expression of Endophilin A1 in DG-75 cells with western blotting. For this reason, we did not further study a role of Endophilin A1. The cortical actin cytoskeleton, however, has been reported to play an important role in BCR clustering and activation in B cells. Specifically, with stimulation, actin has been shown to remodel to allow increased mobilization and clustering of BCRs (Freeman et al., 2011; Hao and August, 2005; Treanor et al., 2011; Treanor et al., 2010). BCR clustering facilitated by actin reorganization is essential for efficient signalosome formation and subsequent cell activation (Harwood and Batista, 2010; Ketchum et al., 2014; Liu et al., 2012). Actin is also involved in the movement of BCR containing vesicles from the early to late endosome after internalization (Brown and Song, 2001). In light of the enhanced recruitment of actin we observed in Supplemental Figure 4, and previous work suggesting a role for actin in BCR clustering and activation, we decided to further investigate the role of actin in BCR endocytosis at these novel clathrin on SRM structures. We likewise pursued the role of FCH and double SH3 domain containing protein 2 (FCHSD2). This protein has been shown to be involved in generating f-actin at sites of classical clathrin mediated endocytosis, is found in B cells, and has been associated as a risk factor in autoimmune diseases (Almeida-Souza et al., 2018; Lessard et al., 2016).

We analyzed actin colocalization with the BCR over 15 minutes in DG-75 cells stimulated with either 2 or 8 ug/mL F(ab)’2 (Figure 5a - c). Cells stimulated with 2 μg/mL F(ab)’2 did not show significant colocalization of the BCR with actin until the late 15 minute time point. However, stimulation with 8 μg/mL F(ab)’2 induced colocalization of BCRs and actin as early as 5 minutes, and this colocalization continued to increase until the 15 minute time point. Thus, with strong antigen simulation, actin appears to be recruited to large BCR clusters early. This suggests a role for actin in endocytosis of large BCR clusters that is different from its role in classical clathrin mediated endocytosis. Figure 5c shows representative images of cells expressing the BCR fused to GFP and stained with phalloidin to mark filamentous actin. Of note, we observed that in unstimulated cells, or cells stimulated with 2 μg/mL F(ab’)2, actin filaments are mostly arranged around the footprint of the cell (Figure 5c, top 3 rows). After stimulation with 8 μg/mL F(ab’)2 for 15 minutes, however, actin appears to reorganize and move away from the footprint and assemble around BCR clusters at the plasma membrane (Figure 5c, bottom row).

**Figure 5.**
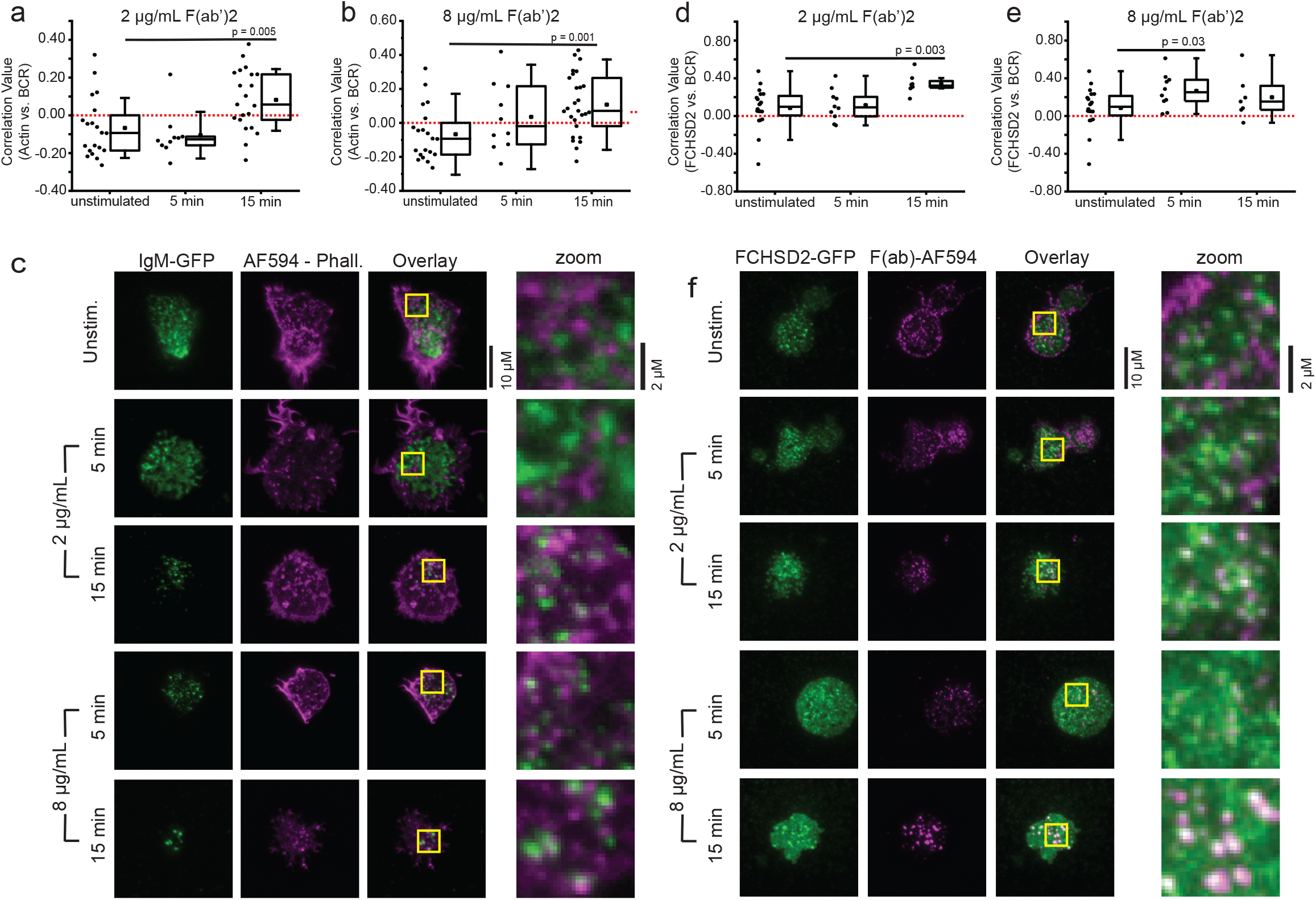
Actin is recruited early to BCR clusters stimulated with a high concentration of F(ab’)2. (a) colocalization analysis of the BCR (IgM-GFP overexpression) and actin (stained with AlexaFluor 594-phalloidin) in DG-75 cells stimulated with 2 μg/mL F(ab’)2 for 5 or 15 minutes. b) the same colocalization analysis done in part a is shown for cells treated with 8 μg/mL F(ab’)2 for 5 or 15 minutes. (c) representative TIRF microscopy images from the colocalization analysis presented in parts a and b. The zoomed images are of from the regions highlighted in yellow boxes. (d) colocalization analysis of the BCR (labeled with AlexaFluor594-F(ab) fragment) and FCHSD2 (FCHSD2-GFP overexpression) in DG-75 cells stimulated with 2 μg/mL F(ab’)2 for 5 or 15 minutes. (e) the same colocalization analysis done in part d is shown for cells treated with 8 μg/mL F(ab’)2. (f) representative TIRF microscopy images from the colocalization analysis presented in parts d and e. All of the box plots from parts a, b, d, and e show the mean (square), median (line), 25/75 percentile range (box), and outliers with a coefficient value of 1.5 and data points (circles) from each cell.

Next, we analyzed the localization of FCHSD2 in relation to clathrin and BCRs. Consistent with previous reports, FCHSD2 is highly colocalized with clathrin (Supplementary Figure 5). We also observed colocalization of FCHSD2 with BCR clusters generated by 2 or 8 μg/mL F(ab’)2 (Figure 5 d-f). Figure 5 f shows representative images of cells expressing FCHSD2-fused to GFP and stained with anti-IgM F(ab) to mark the localization of the BCR. At the 5 minute time point, FCHSD2 is more highly correlated with BCR clusters stimulated with a 8 μg/mL rather than 2 μg/mL. These data are consistent with actin and actin polymerization machinery being specifically recruited to large BCR clusters.

The above data show that actin and FCHSD2 have distinct recruitment kinetics in cells stimulated with 8 μg/mL F(ab’)2. Both proteins are recruited to BCR clusters stimulated with 8 μg/mL F(ab’)2 at an early time point. This is not due to a differential recruitment of clathrin, because clathrin is recruited to BCR clusters with similar kinetics at 2 or 8 μg/mL F(ab’)2 (Supplemental Figure 6). This differential recruitment suggests that both actin and FCHSD2 have distinct functional roles in endocytosis of large BCR clusters. Based on these data, we hypothesize that clathrin and FCHSD2 act together on SRM structures to activate actin polymerization and endocytosis at the early 5 minute time point. While we show that FCHSD2 clearly co-localizes with clathin, future perturbation studies on the role of FCHSD2 in BCR endocytosis are needed to prove a direct mechanistic role for FCHSD2 in polymerizing actin at SRMs.

Next, to gain a more detailed view of the interaction between actin and the BCR cluster in stimulated cells, we used 3D structured illumination microscopy to obtain a sub-diffraction level view of how actin interacts with BCR clusters (Figure 6 a and b). After 5 minutes of treatment with 8 μg/mL F(ab’)2, we observed actin filaments making close contact with many developing BCR clusters (Figure 6 a, yellow arrows). At the 15 minute time point (Figure 6b), we observed actin filaments reorganize around larger BCR clusters to form basket like structures (yellow arrows) that surround the receptors.

**Figure 6.**
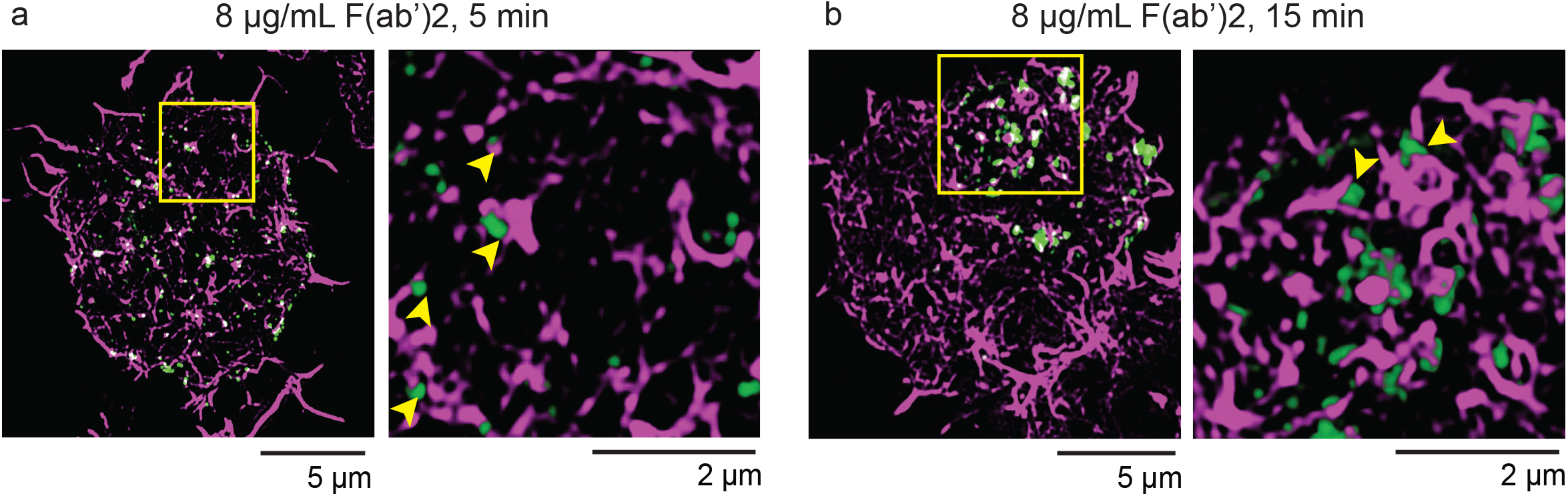
Super-resolution images of actin interaction with BCR clusters. (a and b) 3D Structured Illumination microscopy of cells treated with 8 μg/mL F(ab’)2 for 5 or 15 minutes. DG-75 cells were transfected to express IgM-GFP and then stained with Alexa Fluor 594-Phalloidin after stimulation. Zoomed images correspond to regions in yellow boxes. Yellow arrows in part g show early points of contact between actin and the BCR clusters. The yellow arrows in part h show basket like actin structures formed around BCR clusters.

Together these data show that significant changes in the organization of actin occur after simulation with a high concentration of F(ab’)2. Specifically, BCR clusters colocalize with the f-actin activating protein FCHSD2 and actin (Figure 5) and forms basket like structures around BCR clusters (Figure 6). We hypothesize that structural changes in the localization and arrangement of actin are essential for the formation of smooth raised structure on the membrane capped with clathrin. Next, we tested this hypothesis by perturbing the formation of filamentous actin using the actin disrupting drug Cytochalasin D.

### Actin filaments are essential for efficient formation of clathrin on SRM structures and internalization of large BCR clusters

To perturb the cortical cytoskeleton we used the f-actin destabilizing drug Cytochalasin D to determine what stage of large BCR cluster internalization requires actin: 1) receptor clustering, or 2) formation of clathrin on SRM structures or 3) internalization. We analyzed the effects on receptor clustering using direct STORM localization microscopy to study the BCR. We measured the cluster sizes of unstimulated DG-75 cells or cells stimulated with 8 μg/mL F(ab’)2 in the presence or absence of Cytochalasin D (Figure 7a). As expected, in unstimulated cells there were few large BCR clusters. On average only about 1% of total fluorescence in unstimulated cells was in large clusters. In contrast, cells treated with 8 μg/mL F(ab’)2 alone or stimulated in the presence of Cytochalasin D generated large BCR clusters at similar increased relative percentages (44% and 45% respectively; Figure 7b).These results are consistent with previous reports that actin depolymerization is needed to increase BCR mobilization at the plasma membrane and promote receptor clustering (Freeman et al., 2011; Hao and August, 2005; Treanor et al., 2011; Treanor et al., 2010). Likewise, these data indicate that filamentous actin is not necessary to drive BCR clustering.

**Figure 7.**
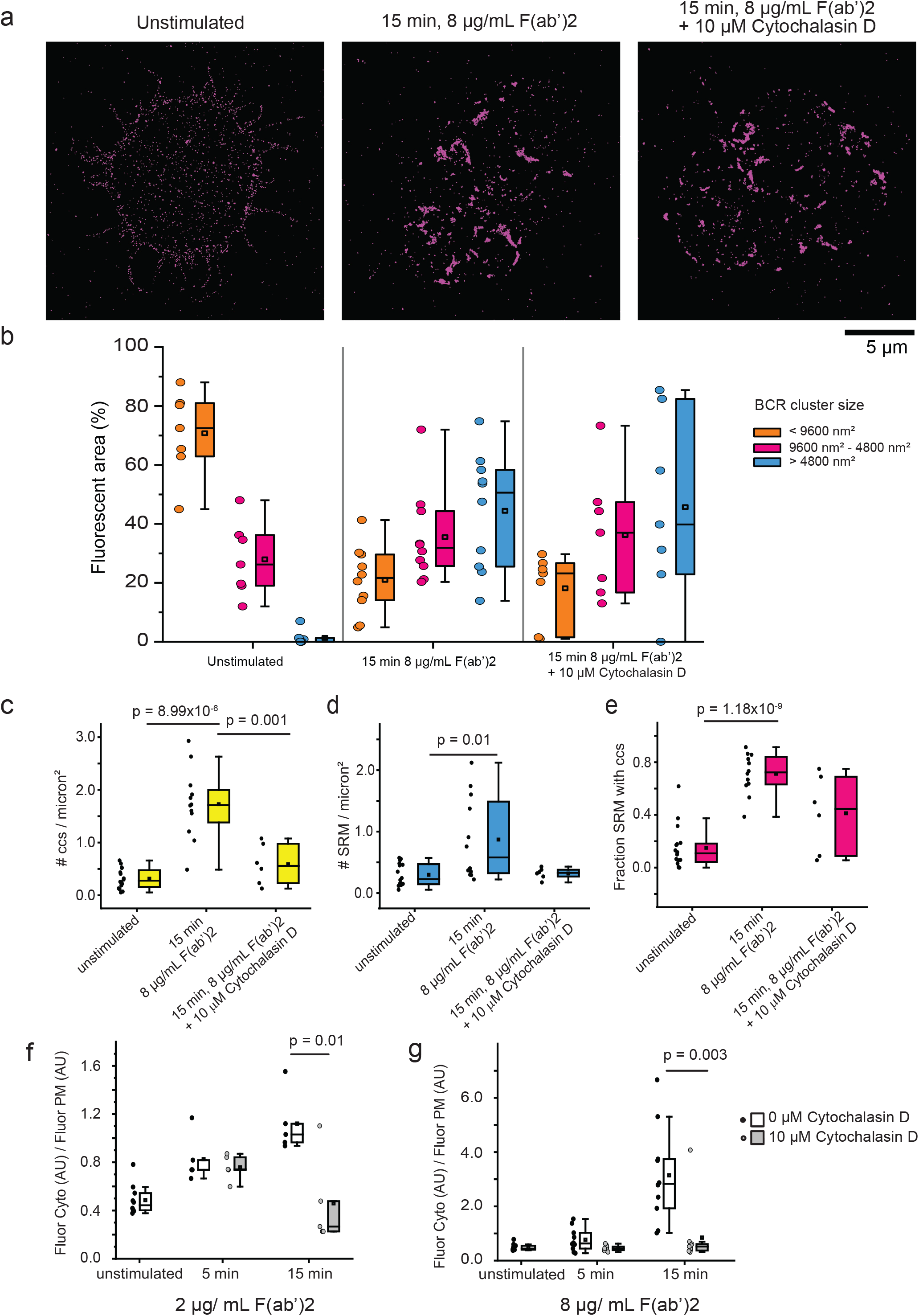
Cytochalasin D blocks efficient biogenesis and internalization of clathin on SRM structures. (a) representative reconstructed dSTORM images of BCR clustering on the bottom surface of DG-75 cells labeled with anti-human IgM F(ab)-Alexa Fluor 647 (magenta) and left unstimulated, stimulated with 8 μg/mL F(ab’)2 alone or stimulated with 8 μg/mL F(ab’)2 in the presence of Cytochalasin D for 15 min prior to plating, fixation, and imaging. (b) Percent of the total fluorescent area composed of small (<9600 nm2, orange bar), intermediate (9600-48000 nm2, pink bar), and large (> 48000 nm2, blue bar) puncta for cells stimulated under the conditions shown in part a (n = unstimulated 7 cells – 15,050 spots, 15 min 8 μg/ml 10 cells – 5,676 spots, 15 min 8 μg/ml + Cytochalasin D 7 cells – 3,914 spots). For parts c-e, cells were stimulated as indicated and then unroofed, and processed by PREM before TEM imaging. TEM images were segmented as previously described for Figure 3, and the density of clathrin (c) and SRM structures (d) is shown. Part (e) shows the fraction of SRM structures that have clathrin on them. (representative masks used for generating these data are shown in Supplemental Figure 6) (f-g)Cells were stimulated with (f) 2 μg/mL or (g) 8 μg/mL F(ab’) 2 alone (open circles and box) or in the presence of 10 μM Cytochalasin D (closed black circles and grey box) before fixation and imaging with a confocal microscope. BCR internalization is presented as the relative fluorescence in the cytoplasm divided by the fluorescence identified at the plasma membrane in a central z-slice for each cell. All Box and whisker plots in this figure show the mean (square), median (line), 25/75 percentile range (box), and outliers with a coefficient value of 1.5.

We next used PREM to analyze the effect of Cytochalasin D on the structure and density of clathrin and SRMs at the plasma membrane (Figure 7c - e). TEM images of membranes from cells that were left untreated, stimulated with 8 μg/mL F(ab’)2 alone or stimulated in the presence of Cytochalasin D were segmented to identify clathrin and smooth raised membrane structures (representative images of clathrin and SRM segmentation are presented in Supplemental Figure 7a - d). In stimulated cells, the density of both clathrin and SRM structures increased compared to untreated cells, and this effect was blocked by Cytochalasin D treatment (Figure 7c and d). Consistent with these results, the fraction of SRM structures with clathrin associated also decreased when cells were treated with Cytochalasin D, indicating a partial inhibitory effect of Cytochalasin D on clathrin/SRM structure biogenesis (Figure 7e). Interestingly, some clathrin on SRM structures were still present in cells treated with Cytochalasin D. This suggests that additional proteins other than actin are involved in the generation of clathrin/SRM structures or that the block of actin’s actions with Cytochalasin D was partial. We also observed less clustering and a larger average radius of both clathrin and SRM structures in cells treated with Cytochalasin D (Supplementary Figure 7e-h).

Finally, we tested the effect of cytochalasin D on internalization of BCRs with a fluorescent internalization assay (Figure 7f and g). At low concentration of F(ab’)2 stimulation, BCR internalization was inhibited after 15 minutes of stimulation (Figure 7f). In contrast, cells stimulated with a high concentration of F(ab’)2 were sensitive to Cytochalasin D treatment at the earlier 5 minute time point (Figure 7g). The robust and rapid inhibition of BCR internalization into the cytosol of cells stimulated with high concentrations of ligand suggests that actin is essential for late steps in endocytosis. Together, our data indicate that actin has a complex mechanistic role in the formation and internalization of these specialized endocytic structures in B cells. Specifically, actin is preferentially recruited at an early time point in cells stimulated with high concentrations of ligand and is required for the efficient biogenesis of clathrin on SRM structures. Furthermore, the almost complete block of internalization indicates that either fission or transport of large BCR clusters into the cytosol is actin dependent.

## DISCUSSION

Adaptive immunity depends on the ability of B cells to capture antigen through a regulated internalization mechanism of the B cell receptor (Harwood and Batista, 2010; Hoogeboom and Tolar, 2016). Here, we provide direct structural evidence that the mechanism of BCR capture changes with the concentration of the stimulating ligand and the resulting changes in size of BCR clusters. In unstimulated cells, BCR is evenly distributed across the plasma membrane. The plasma membrane in these cells contains few clathrin-coated structures compared to other mammalian cell types. Upon antigen stimulation at low concentration, BCRs cluster and associate with an increased number of classical clathrin coated structures which are likely able to internalize these small BCR clusters using the standard clathrin mediated endocytosis pathway. At higher antigen concentrations and longer times, however, large clathrin-associated BCR clusters coalesce onto smooth raised membrane invaginations of the plasma membrane. FCHSD2 and actin colocalize with these large BCR clusters within minutes and forms cup-like structures around the clusters. Organized actin is required for internalization of these large BCR clusters into the cytosol. Our data support a model where B cells can accommodate the internalization of a wide range of BCR cluster sizes by deploying either classic clathrin mediated endocytosis and a hybrid form of endocytosis that uses clathrin coats to gather receptors onto large membrane carriers to capture receptors wholesale into the cytoplasm. The latter is distinctly sensitive to actin disruption at an early time point.

The role of actin in the formation and internalization of large BCR clusters is complex. Our data suggests that actin plays two mechanistic roles. First, at the early time points actin is recruited to BCR clusters, possibly by the FCHSD2 protein. FCHSD2 has been shown in other studies to activate actin polymerization (Almeida-Souza et al., 2018). At this stage actin plays a role in supporting increased clathrin recruitment to the plasma membrane and in the formation of smooth raised membrane structures (Figure 7 c and d). At later stages, actin is essential for internalization of smooth raised membrane structures into the cytosol (Figure 7 g). Although this dual role of actin in endocytosis of large BCR clusters was unexpected, it is not unprecedented for endocytic proteins to play more than one role in caputring surface receptors. For example, the GTPase Dynamin has been extensively studied for its role in driving membrane scission at late stages of clathrin mediated endocytosis (Mettlen et al., 2018). Specifically, the two isoforms of Dynamin (Dyn1 and Dyn2) play unique roles at the early and late stages of clathrin mediated endocytosis (Srinivasan et al., 2018). Dyn1 controls the early initiation of clathrin coated pits while Dyn2 drives membrane scission at the late stage of clathrin mediated endocytosis.

We consider several implications of our finding on the B cell endocytic machinery. First, in unstimulated cells most BCR was not associated with clathrin. Nor were there many clathrin structures present at the membrane. Most mammalian cells maintain a dynamic population of clathrin-coated structures at the membrane at a density between 0.5 and 2 structures per square micron (Dambournet et al., 2018; Sochacki et al., 2012; Taylor et al., 2011). These structures support constitutive endocytosis and the recycling of many key receptors such as the transferrin receptor. We find that resting B cells maintain very few clathrin structures at their plasma membrane. This could serve to increase the steady-state concentration of BCRs on the surface. In contrast to unstimulated cells, most BCR fluorescence was associated with clathrin structures in cells treated with 8 μg/mL F(ab’)2 for 15 minutes. The relative absence of BCR-negative clathrin coated pits suggests that clathrin assembles specifically on antigen-induced BCR clusters. Considering the need for immune responses to be specific, regulated, and rapid, the lack of membrane-associated clathrin structures in resting B cells could serve as a barrier to limit BCR endocytosis and minimize off-target or autoimmune responses that might occur as a result of nonspecific antigen internalization (Davis et al., 2010). Furthermore, the selective recruitment of clathrin to clustered BCR would ensure specific and efficient internalization of the target antigen.

Second, clathrin-associated BCR clusters form and group together in a concentration-dependent manner following F(ab’)2 stimulation. Treatment with 2 μg/mL F(ab’)2 stimulates efficient BCR internalization by flow cytometry, and this internalization appears to involve clathrin, based on colocalization of BCR with clathrin in confocal microscopy images (Supplementary Figure 1e). However, CLEM data showed that this does not yield dramatic growth of BCR clusters or recruit large amounts of additional clathrin to the plasma membrane. In contrast, after 5 minutes of stimulation with 8 μg/mL F(ab’)2, the amount of clathrin at the surface membrane increases and nearly all BCR clusters are associated with clathrin. After 15 minutes of stimulation with 8 μg/mL F(ab’)2, BCR clusters capped with clathrin increase in number, cluster together, and are found to be largely associated with smooth, raised membrane structures. Thus, individual clathrin structures precede the development of SRMs capped with clathrin and BCRs. These data suggest a model in which relatively small BCR clusters may be internalized through a classic clathrin-mediated endocytic pathway, and BCR clusters that grow too large to be contained in a small clathrin-coated vesicle could require a hybrid mechanism of membrane internalization that involves bulk plasma membrane remodeling and early recruitment of actin. Given the large body of literature that supports a role for lipid microdomains in BCR signaling, one avenue for future work is to examine whether the smooth, raised structures we observe by CLEM form as a result of phase transitions within the plasma membrane (Cheng et al., 1999; Sezgin et al., 2017; Stone et al., 2017).

From this model, we suggest that BCR clusters of different sizes would pose distinct challenges for the endocytic machinery and may therefore recruit distinct sets of regulatory proteins in each case. Whereas small BCR clusters may be internalized by a basic set of clathrin-associated proteins, perhaps larger BCR clusters depend on an expanded set of endocytic proteins (Ferreira and Boucrot, 2018; Watanabe and Boucrot, 2017). In support of this theory, we have shown that large BCR clusters rely on increased involvement of actin for internalization. This is analogous to what very large antigens appear to do in a mechanism that more closely resembles phagocytosis (Zhu et al., 2016). Our survey of endocytic proteins in Supplemental Figure 4 also identified some proteins that are differentially recruited to small and large BCR clusters and could be further investigated. Additionally, previous work has shown that B cells rely on myosin to a greater extent when internalizing antigen presented on a bilayer compared to soluble antigen (Natkanski et al., 2013). Similar to these studies, which show context and antigen-dependent variability in the mechanism of BCR uptake, our data support the hypothesis that BCR cluster composition and size may be another key factor that impacts the mechanism of BCR internalization in B cells.

We have shown that endocytic structures associated with large BCR clusters exhibit a unique nanoscale morphology at the plasma membrane. These structures exhibit dense closely-packed arrays of small clathrin structures on smooth, raised membrane invaginations. These clathrin-coated sites are unlike those observed by EM in other mammalian cell types (Dambournet et al., 2018; Scott et al., 2018; Sochacki et al., 2017; Sochacki et al., 2014). However, there is some similarity between the raised clathrin structures we observe in B cells and the structures that form on mast cells in response to FCeRI clustering (Aggeler and Werb, 1982; Cleyrat et al., 2013; Wilson et al., 2002). There is overlap in the BCR and FCeRI signaling cascades (Siraganian et al., 2010), and the morphological similarity we observe in the receptors’ endocytic structures suggests that there may be commonalities in their mechanisms of internalization as well. Future work will be needed to determine the identity of other molecules that drive and regulate the proteins required for biogenesis of smooth, raised structures at the plasma membrane.

Our data supports the idea that BCR endocytosis occurs by an adaptable mechanism, employing different nanoscale endocytic structures to internalize BCR clusters of different sizes. It remains to be determined whether these distinct endocytic structures traffic differently throughout the cell and what proteins might be employed to specifically drive these invaginations. Future studies to understand whether these structurally distinct endocytic pathways influence the efficiency of antigen processing and presentation will be valuable to ensure that future adjuvants and immunogens are optimally designed to cooperate with the available B cell endocytic machinery.

## MATERIALS AND METHODS

### Cell lines and imaging reagents

The IgM+VkL+ human DG-75 B cell line was purchased from ATCC (CRL-2625) and cultured in a growth medium consisting of RPMI 1640 (−) Phenol Red with 10% FBS, 1% Penicillin/Streptomycin, 10mM HEPES pH 7.4, and 1mM Sodium Pyruvate. Cells were periodically tested for mycoplasma. Cell line identity was verified as a human B cell (ATCC). Anti-human IgM F(ab) and F(ab’)2 fragments were purchased from Jackson Immunoresearch laboratories, including: Goat Anti-Human IgM Fc5μ F(ab)-Fluorescein (109-097-043), Goat Anti-Human IgM Fc5μ F(ab)-Alexa Fluor 647 (109-607-043), Goat Anti-Human IgM Fc5μ F(ab)-Alexa Fluor 488 (109-547-043), Goat Anti-Human IgM Fc5μ F(ab)-Alexa Fluor 594 (109-587-043), and AffiniPure F(ab’)2 Fragment Goat Anti-Human IgG + IgM (H+L) (109-006-127). Fixable Viability Dye eFluor450 (65-0863-18), Alexa Fluor 488 Phalloidin (A12379), mouse anti-clathrin heavy chain antibody clone X-22 (MA1065), and donkey anti-mouse Alexa Fluor647 (A31571) were purchased from ThermoFisher. Coverslips for TIRF and confocal microscopy #1.5 25mm were purchased from Warner Instruments (64-0715), and high density (400 particles /100 mm2) wide spectral band (600+/− 100 nm) gold fiducial coverslips were purchased from Hestzig LLC. for correlative fluorescence electron microscopy. Coverslips were washed in a heated solution of hydrogen peroxide and ammonia solution as previously described (Trexler and Taraska, 2017) with gold fiducial coverslips exposed to the cleaning solution for 10 minutes. Poly-L-Lysine for coating coverslips was purchased from Sigma Aldrich (P4832).

### Construction of DG-75 IgM-GFP

The DG-75 IgM VH sequence [Genbank: Z74668, (Chapman et al., 1996)] with the native VH3-23*01 leader peptide sequence [IMGT: M99660, (Matsuda et al., 1993)],was synthesized by Biobasic and cloned by overlap PCR in frame with the human IGHM sequence obtained from DNASU (HscD00445046). This fragment was then cloned by overlap PCR upstream of a PCR product encoding meGFP with an SGSGSGGSGSGGGPVAT N-terminal linker peptide. PCR reactions were carried out using Accuprime Pfx Supermix (ThermoFisher: 12344040) with primers synthesized by Integrated DNA Technologies. The DG75-VH-IgM-linker-GFP insert was digested with NheI/NotI and ligated into the Clontech pEGFP-N1 vector.

### Flow cytometry-based BCR internalization assay

DG-75 cells at a concentration of 1-1.5 million cells /mL were spun down at 200xg for 5 minutes. The cell pellet was resuspended at 10 million cells/mL in a staining medium of RPMI complete with the addition of 1:1000 anti-human IgM F(ab)-FITC, 1:1000 anti-human IgM-F(ab)-Alexa Fluor 647, and 1:1000 eFluor450 live-dead violet. Cells were stained at 4°C for 30 minutes prior to washing with RPMI complete and pelleting cells at 200xg for 5 minutes. The stained, washed cell pellet was resuspended to a concentration of 2 million cells/mL in RPMI complete, filtered through a 40-μm filter, and 700 μL of cells were aliquoted into each round bottom tube. Anti-human IgM F(ab’)2 was added to a final concentration of 0, 0.8, 1.7, 3.5, 7, or 14 μg/mL immediately prior to flow cytometry analysis and a temperature shift to 37°C. Cells were incubated at 37°C and analyzed every 10 minutes post-F(ab’)2 addition for up to 60 minutes. Gates were drawn to isolate singlet, live (eFluor450 negative) cells.

BCR internalization was measured by tracking the pH-induced decrease in FITC fluorescence that occurs as the BCR-bound FITC-conjugated F(ab) is internalized and trafficked into low pH endosomal compartments. To account for the change in fluorescence that may result from dissociation of fluorophore-conjugated F(ab) from the B cell surface over time, the change in FITC signal was corrected for the measured decrease in the fluorescence of the pH-insensitive fluorophore, Alexa Fluor 647. BCR internalization plots are presented as the negative change in FITC fluorescence relative to that of Alexa Fluor 647, and are calculated as follows:

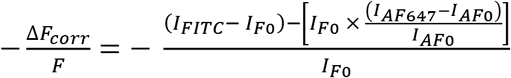

Where *I*_*FITC*_ is the cell population mean fluorescence intensity (MFI) in the FITC channel at time of measurement, *I*_*F0*_ is the initial MFI of the cell population in the FITC channel, *I*_*AF647*_ is the cell population MFI in the Alexa Fluor 647 channel at time of measurement, and *I*_*AF0*_ is the initial MFI of the cell population in the Alexa Fluor 647 channel.

### Live cell confocal laser scanning microscopy

Imaging of BCR clustering and internalization in live cells in three dimensions was carried out using a Zeiss 880 confocal laser scanning microscope with a 63X, 1.4NA oil-immersion PlanApo DIC objective. DG-75 cells were prepared for imaging by spinning down 1 million cells at 200xg for 5 min, and resuspending the pellet in 100 μL of growth media with 1:1000 anti-human IgM-F(ab)-Alexa Fluor 488. Fluorescent labeling of the BCR was carried out at 4°C for 30 minutes. Cells were then diluted with 900 μL of imaging buffer, pelleted at 200xg for 5 min, and resuspended in 150 μL IB. Approximately 300,000 labeled cells were allowed to settle for 5 min onto a cleaned, poly-lysine treated, rinsed, coverslip assembled in an Attofluor chamber (ThermoFisher: A7816) imaging chamber containing 600 μL of imaging buffer on a stage heated to 37°C inside an environmental control box. Regions containing multiple live, healthy cells were identified and the equatorial z-position was set as the origin. Two planes above and below this equatorial position were imaged, with a z-step size of 0.43 μm, encompassing a thickness of ~2.1 μM in the middle of the cell. After the addition of anti-human IgM F(ab’)2 to a final concentration of 12 μg/mL, this five slice stack was imaged once every 6 seconds for a total of 30 min using 488 nm excitation and detecting transmitted and emitted light using T-PMT and GaAsP detectors, respectively.

### Immunofluorescence imaging of BCR, actin, and clathrin

To observe endogenous clathrin relative to BCR on the plasma membrane, DG-75 cells were harvested at a concentration of 1-1.5M cells/mL and spun down at 200xg for 5 minutes. The cell pellet was resuspended at 10 million cells/mL in growth medium containing 1:5000 anti-human IgM F(ab)-Alexa Fluor 594. Cells were labeled for 30 minutes at 4°C followed by a 5-fold dilution in growth medium and centrifugation at 200xg for 5 minutes. The labeled cell pellet was resuspended in growth medium to achieve a density of 2 million cells/mL, and anti-human IgM F(ab’)2 was added to achieve a F(ab’)2 concentration of 2 or 8 μg/mL (or omitted for unstimulated cells). F(ab’)2-stimulated cells were incubated at 37°C for 5 or 15 minutes before diluting the cells 2-fold with cold IB. Approximately 125,000 cells were allowed to settle for 12 minutes onto cleaned, poly-lysine treated, rinsed coverslips in a six-well plate containing 2 mL IB per well. Coverslips were treated with 0.5% paraformaldehyde (PFA) (Electron Microscopy Sciences) in IB for 1 minute before transferring them into 2% PFA in imaging buffer for 20 minutes. Fixed coverslips were washed with PBS, permeabilized by the addition of a solution of 0.5% TritonX-100 and 3% BSA in PBS for 2 minutes, and blocked in a solution of 0.2% TritonX-100 and 3% BSA in PBS for 1 hour. Coverslips were incubated in primary mouse anti-clathrin heavy chain antibody X-22 (6 μg/mL in blocking buffer) for 1 hour, washed with blocking buffer, and incubated with secondary donkey anti-mouse-Alexa Fluor 647 antibody (2 μg/mL) and phalloidin-Alexa Fluor 488 (33 nM) in blocking buffer for 1 hour. After washing in blocking buffer and PBS, coverslips were fixed for 10 minutes in 2% PFA in PBS, rinsed in PBS, and stored at 4°C protected from light until imaging. Fixed, immunostained cells were imaged on the same Zeiss 880 confocal laser scanning microscope setup used for live cell imaging, using a 3-track program with excitation at 488, 561, and 633 nm. Z-stacks were collected to encompass the full cell volume.

### Direct stochastic optical reconstruction microscopy (dSTORM)

The super-resolution localization method, dSTORM, was used to measure nanoscale changes in the size of BCR clusters on the plasma membrane following stimulation with anti-human IgM F(ab’)2. DG-75 cells were prepared for imaging exactly as described for immunofluorescence imaging, but labeled with anti-human IgM F(ab)-Alexa Fluor 647 (1:5000 in growth medium) and without permeabilization or additional staining for clathrin or actin. Fixed, PBS washed coverslips were assembled in sealed AttoFluor imaging chambers (Thermo Fisher) containing a freshly-prepared dSTORM buffer (10% w/v D-glucose, 100 mM beta-mercaptoethanol, 0.8 mg/mL glucose oxidase, and 0.04 mg/mL catalase in PBS). Imaging was carried out as previously described in detail using a Nikon Eclipse Ti inverted microscope with a 100X, 1.49NA objective, and an Andor iXon Ultra 897 EM-CCD camera under the control of Nikon Elements NSTORM software (Sochacki and Taraska, 2017). STORM imaging was performed in TIRF, with 647 nm excitation at 75% laser power, collecting 10-20,000 frames with 10 msec exposure times. STORM images were processed as described previously, with the following modifications: a minimum height of 200 was used in the identification parameters for all images, and a value of 200 was used as the minimum number of photons during filtering. Drift correction was performed in Nikon Elements software, and Tiff images were rendered using a cross representation with 8 nm pixel size. Localization precision was measured using fourier shell correlation and measurement of individual localizations in the rendered Tiff image files, both of which yielded an apparent precision of ~27 nm.

In order to measure BCR cluster sizes from the STORM data, images were maximum filtered using a radius of 1 pixel (8 nm), and thresholded using the Li filter in Fiji/ImageJ (Schindelin et al., 2012) to produce a down-sampled, binary image, with a resolution of 40 nm. The result of this processing is to cluster localizations that are separated in space by a distance of less than the precision of the original image, 24 nm, edge-to-edge – a conservative distance cutoff, which was empirically determined to identify reasonable clusters across the conditions tested i.e. monodisperse, small punctae in unstimulated cells, and heterogeneous, larger cluster sizes in stimulated cells. Six to twelve cells were analyzed for each condition, and the areas of all particles in the thresholded images were measured in Fiji/ImageJ. The total fluorescent area was measured for each cell, and the percent of this area composed of small (<9600 nm^2^), intermediate (9600 – 48000 nm^2^), and large (>48000 nm^2^) fluorescent punctae was plotted in Origin (OriginLab). As described in the results section, these area cutoffs were selected based on the typical size of clathrin coated pits.

### Correlative super-resolution fluorescence and electron microscopy

To directly visualize endocytic structures associated with BCR fluorescence, DG-75 cells were prepared for imaging exactly as described for STORM imaging above, with a few modifications. After 12 minutes of attachment to poly-L-lysine coated gold fiducial coverslips (Hestzig, LLC) cells were fixed and ‘unroofed’ to expose the cytoplasmic surface of the membrane adhered to the coverslip by squirting 4 mL of 0.75% PFA in stabilization buffer (70 mM KCl, 5 mM MgCl_2_, 30 mM HEPES, pH 7.4) steadily onto each coverslip using a 10-mL syringe with a 21Gx1.5 needle, moving the stream over the coverslip at a right angle. After unroofing, coverslips were fixed in 2% PFA in stabilization buffer for 20 minutes, rinsed with PBS, treated with phalloidin-Alexa Fluor 488 (33 nM in PBS) for 10 minutes, and rinsed again in PBS prior to imaging.

Labeled, unroofed coverslips were assembled in sealed AttoFluor chambers containing freshly prepared dSTORM buffer. Imaging was performed as described above, with some additional steps to map cells on the coverslip for later correlation with TEM images. As described elsewhere (Sochacki et al., 2017), a ~1mm^2^ region of the coverslip was initially mapped by montaging a 15×15 area in epifluorescence with excitation at 488 and 647 nm. These large montages of the phalloidin-Alexa Fluor 488 label were useful for re-identification of the cells imaged by STORM prior to imaging platinum replicas by TEM. After imaging 5-8 cells, a 4 mm diameter circle was etched on the bottom of the coverslip with a diamond objective (Leica) to indicate the area of interest during sample processing for EM. Etched coverslips were cleaned of oil and fixed in a 2% solution of glutaraldehyde in PBS (Electron Microscopy Sciences).

As described previously (Sochacki et al., 2017) coverslips were prepared for EM by treatment with 0.1% tannic acid solution for 20 min, rinsing with water, and staining with 0.1% uranyl acetate for 20 minutes. Coverslips were dehydrated using a Tousimis critical point dryer, and coated with platinum and carbon in a JEOL freeze fracture device (Sochacki et al., 2012). After locating cells of interest under a 10X phase-contrast objective, carbon-platinum replicas were separated from the underlying glass coverslip using 5% hydrofluoric acid, and lifted onto glow-discharged formvar-coated 75 mesh copper grids for imaging on a JEOL 1400 TEM using SerialEM software for montaging (Mastronarde, 2005), at a magnification of 15,000x.

STORM images were processed as described above, but Tiff images were rendered using a Gaussian representation with 8 nm pixel size. STORM image files were transformed using custom MATLAB software to align images based on the measured coordinates of gold nanorods visible in both the rendered dSTORM Tiff files and TEM montages.

### DG-75 plasma membrane ferritin labeling

49 mg of cationized ferritin from horse spleen (Sigma-Aldrich F1897) was sonicated for 10-15 seconds and then added to three million DG-75 cells for each condition. Samples were then gently swirled by hand to disperse ferritin. Next, the samples were incubated on ice for 10 minutes to allow the ferritin to adhere to cell membranes and then warmed at 37°C for 30 seconds. The samples were spun in a mini-centrifuge in 10 second interval bursts to produce a pellet of approximately 2 mm × 3 mm. The unstimulated sample was then immediately fixed with 2.5% glutaraldehyde and 1% paraformaldehyde in 0.12 M sodium cacodylate buffer at pH 7.2 for 1 hour at room temperature. The stimulated sample was resuspended in PBS and administered F(ab’)2 at a final concentration of 8 μg/ml and incubated for 15 minutes at 37°C. After incubation, stimulated cells were fixed in the same manner described for the unstimulated sample.

### Thin-section transmission electron microscopy

Fixed samples were washed 3 times (20 minutes each) with 0.12 M sodium cacodylate buffer and stained with 1% osmium tetroxide (on ice and in the dark) for 1 hour, washed twice (10 minutes each) in water, and stained with 1% uranyl acetate (overnight at 4°C). The following day samples were processed through a dehydration protocol of increasing concentrations of ethanol, infiltrated with Epon resin, and polymerized at 60°C for at least 48 hours (VWR Symphony oven). Post-staining was determined to be not necessary due to sufficient contrast observed for ferritin. 65-70 nm ultrathin sections were obtained on a Leica-Reichert Ultracut microtome with a diamond knife (Delaware Diamond Knives, Inc.) after each sample block resin was trimmed by a glass knife, and confirmed to be in the cell layer through thick sectioning and Toluidene blue dye. Ultrathin sections were imaged on 200 mesh grids in a JEOL 1400 TEM.

### Analysis of EM and CLEM images

Platinum replica EM images of the entire visible plasma membrane of each cell were segmented by hand using ImageJ/FIJI with ROIs corresponding to the 1) total plasma membrane area, 2) clathrin-coated structures, 3) smooth-raised membranes, and 4) gold fiducial particles. These ROIs were used to calculate density of structures, area of individual structures, Feret diameter of single structures, and nearest-neighbor distances between structures. Analysis, statistics, and plotting were done in Origin (OriginLab). The overlap pixel area between clathrin structures and SRMs was calculated from image masks made from ROIs in Matlab (MathWorks).

The distribution and density of super-resolution dSTORM fluorescence signal excluding regions 0.5 micron radius from gold fiducials were analyzed using the above EM identified clathrin/SRM segmentation masks. Fluorescence signal within and surrounding a 200 nm radius of each segmented membrane structure was measured against the total fluorescence in the surrounding 1.8 micron radius area using Matlab. The resulting fluorescence ratio measurements for each structure in all experimental conditions were plotted in Origin (Originlab).

### Structured Illumination Microscopy

Three Dimensional structured illumination images were collected on a DeltaVision OMX SR microscope with a 60x 1.42NA oil emersion PSF objective. DG75 cells were transfected and stimulated as previously described and then fixed in 4% paraformaldehyde for 15 minutes. For staining with Alexa Fluor 594 Phalloidin, the cells were additionally permeabilized with 0.1% Triton X-100 for 5 minutes, washed with Phosphate Buffered Saline, and then incubated with Alexa Fluor 594 Phalloidin diluted 1:50 in 1% BSA for 30 minutes. For 3D SIM image collection, z-stacks were taken at a focal plane near the cell membrane with a 0.125 μm z-step for a total of about 1.125 μm. Processed z-stacks collected in this way were then reconstructed in Image J using the 3D-Script plugin.

## AUTHOR CONTRIBUTIONS

ADR, TMD and JWT designed experiments. TMD performed molecular biology, confocal, FACS, and TIRF experiments for initial characterization of the system. ADR performed molecular biology, confocal internalization assays, TIRF, SIM, and STORM experiments characterizing the role of actin in BCR stimulation. TMD, AMD, and KAS performed platinum replica TEM and CLEM experiments. KAS designed and wrote analysis software. RA and ADR did thin section TEM. ADR, TMD and JWT wrote the manuscript and all authors commented on work. JWT oversaw the project.

## ACKNOWLEDGMENTS

We would like to thank the NHLBI Electron Microscopy core for support with EM imaging and instrumentation, Xufeng Wu and the NHLBI light Microscopy core for support with fluorescence imaging and instrumentation, the NHLBI FACS core for support with flow cytometry, and all members of the Taraska lab for comments and scientific dialogue. We would like to thank Agila Somasundaram for critically reading the manuscript. We would like to acknowledge Dr. Avital Rodal and Dr. Shiyu Wang for generously sending us plasmids with the FCHSD2-GFP gene. JWT is supported by the Intramural Research Program, National Heart Lung and Blood Institute, National Institutes of Health, Bethesda, Maryland.

---

**Supplementary Figure 1.**
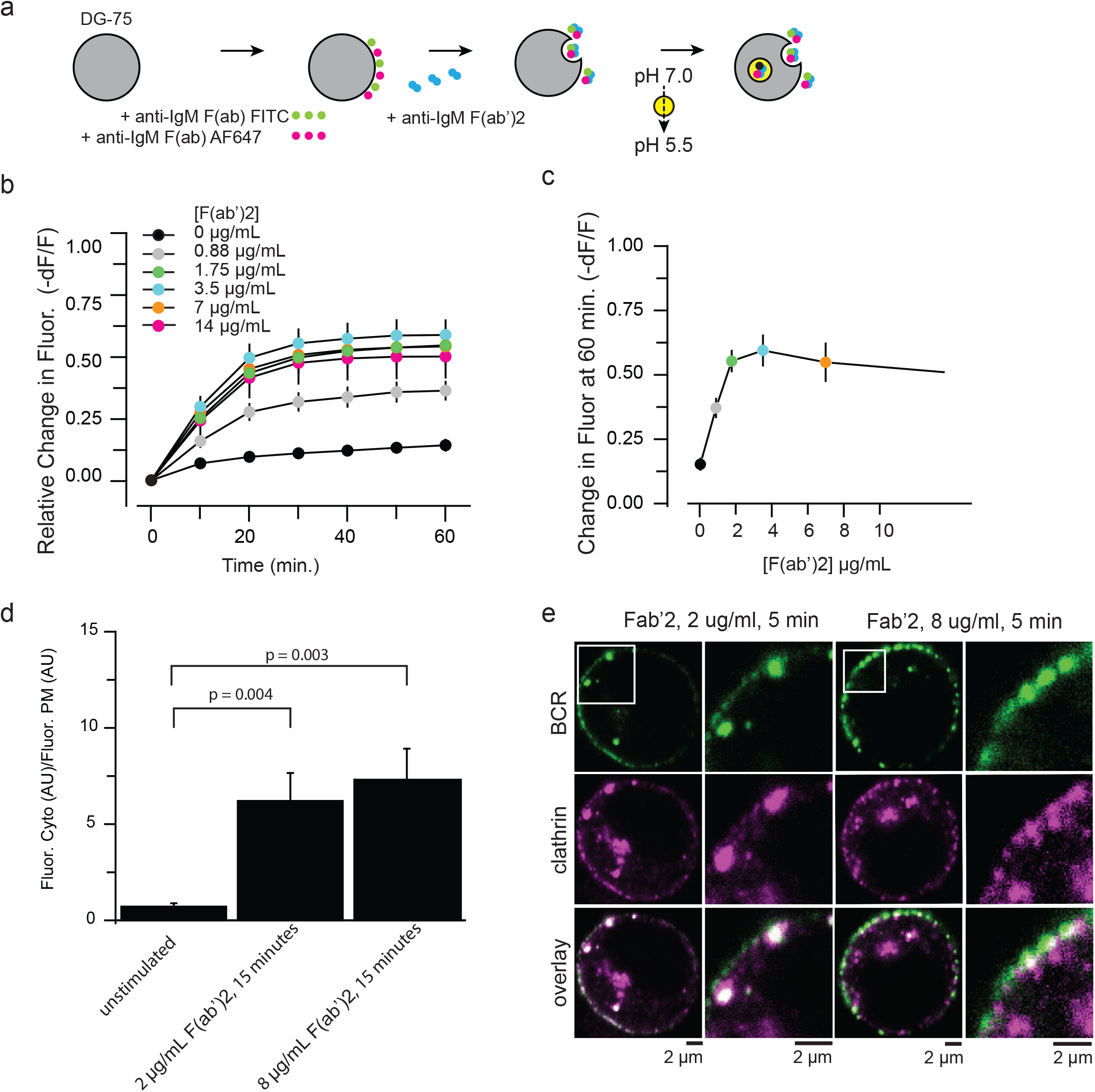
FACS analysis of BCR internalization in human DG-75 cells. (a) Diagram of the flow-cytometry assay developed to measure BCR internalization in DG-75 cells following the addition of F(ab’)2. (b) Plot of BCR internalization measured by the change in fluorescence of anti-human IgM F(ab) FITC relative to the change in fluorescence of anti-human IgM F(ab) Alexa Fluor 647 over time following the addition of anti-human IgM F(ab’)2 at concentrations ranging from 0-14 μg/mL. (c) Plot of BCR internalization at 60 min of stimulation as a function of varying concentrations of anti-human IgM F(ab’)2. Error in (b) and (c) indicate standard deviation across five experiments. (d) Plot of F(ab’)2-induced uptake of fluorescent IgM F(ab)s measured in equatorial confocal sections. Total fluorescence of the plasma membrane compared to the total fluorescence in the cytosol for three experimental conditions to measure BCR uptake as a function of antigen concentration and time (unstimulated, n = 8cells; 2 μg/ml 15 min, 12 cells; 8 μg/ml, 11 cells). The plasma membrane region was segmented according to the ring-like actin cortex stating. Statistical analysis to compare treatment means was a two-sample t-test. P-values between samples are indicated on bar graph. Error bars are SEM. (e) Colocalization of BCR and clathrin in immunofluorescence confocal images of DG-75 cells. Cells were labeled with anti-human IgM Fab-AF488 (green) and stimulated with 2 or 8 μg/mL of Fab’2 for 5 minutes prior fixation and staining for clathrin (magenta).

**Supplementary Figure 2.**
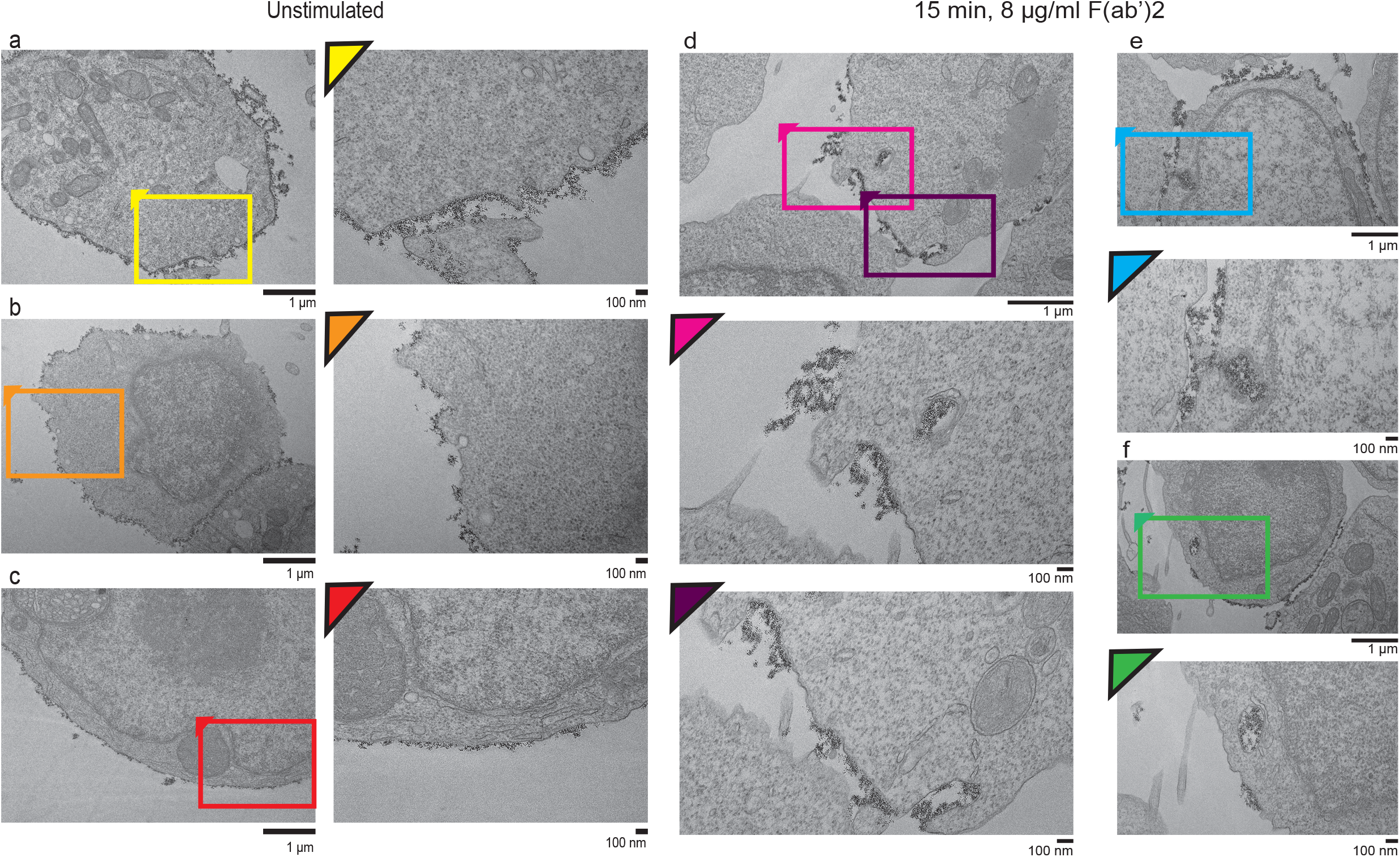
TEM analysis of the resting B cell plasma membrane and membrane morphological changes during BCR clustering 15 minutes post F(ab’)2 stimulation. (a-c) Representative thin-section TEM images of DG-75 cells treated with extracellular cationized ferritin (15 nm black dots) to label the outer leaflet of the plasma membrane and the inner leaflet of endocytic carriers. Zoomed regions (colored boxes) shown on the right side of each larger image show details of membrane morphology. (d-f) Representative thin-section TEM image of DG-75 cells treated with extracellular cationic ferritin and 8 μg/mL F(ab’)2 for 15 minutes to activate and cluster BCRs. Two zoomed regions (colored boxes) from the same cell shown in part d show details of membrane morphology of large endocytic carriers that take up ferritin. Two additional stimulated cells and zoomed regions (colored boxes) are shown in e-f.

**Supplemental Figure 3.**
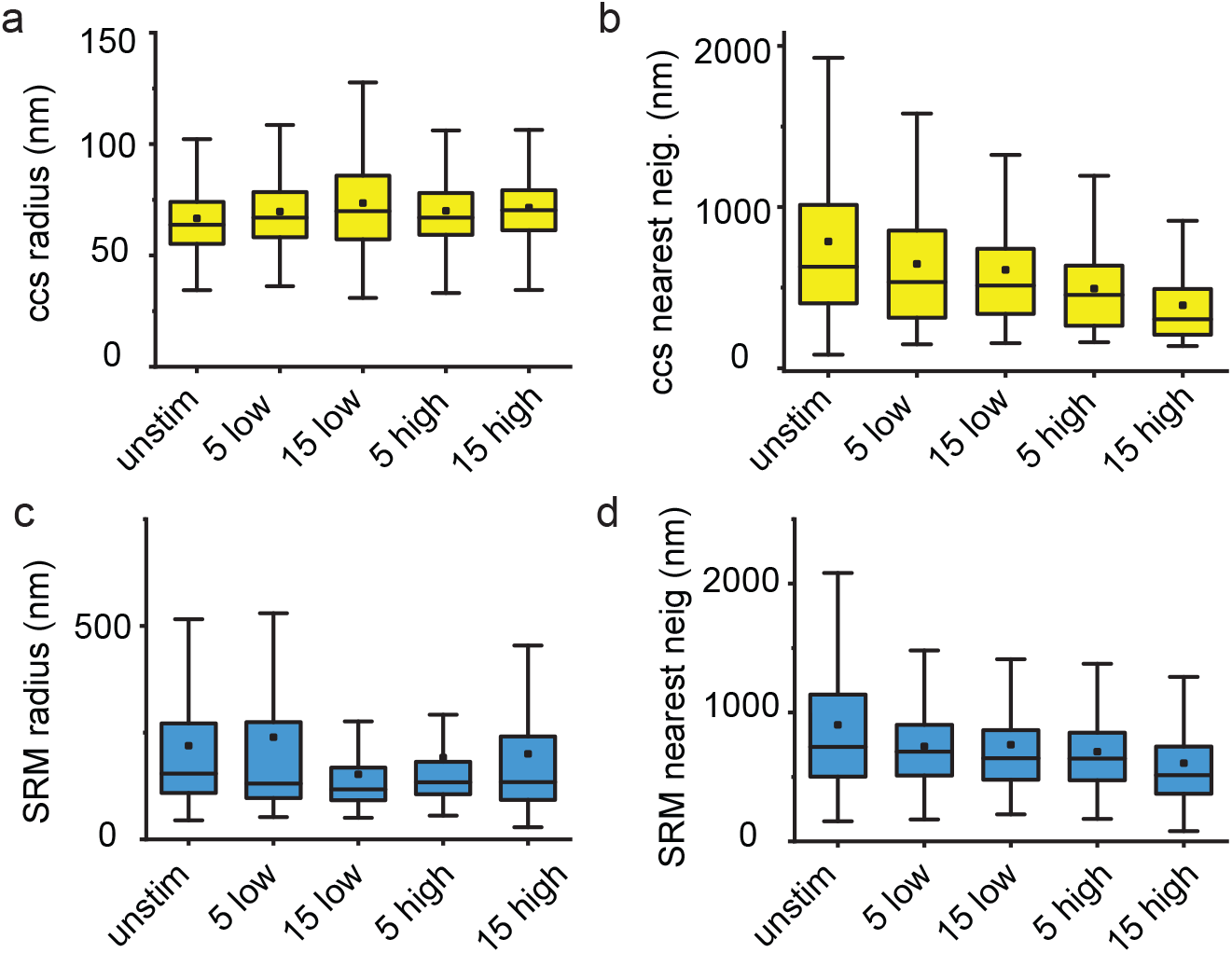
CCS and SRM radius and nearest neighbor distances following F(ab’)2 stimulation. The segmented masks shown in Figure 5 were used to measure changes in CCS radius(a), and (b) the nearest-neighbor distances between CCS (n = Unstim. 16 cells, 820 ccs structures, 5 min 2 μg/ml 9 cells, 486 ccs structures, 5 min 8 μg/ml 6 cells, 482 ccs structures, 15 min 2 μg/ml 6 cells, 499 ccs structures, 15 min 8 μg/ml 12 cells, 2063 ccs structures). Masks were also used to measure SRM radius (c), and nearest-neighbor distance (d) (n = Unstim. 16 cells, 708 SRM structures, 5 min 2 μg/ml 9 cells, 553 SRM structures, 5 min 8 μg/ml 6 cells, 313 SRM structures, 15 min 2 μg/ml 6 cells, 368 SRM structures, 15 min 8 μg/ml 12 cells, 870 SRM structures).

**Supplemental Figure 4.**
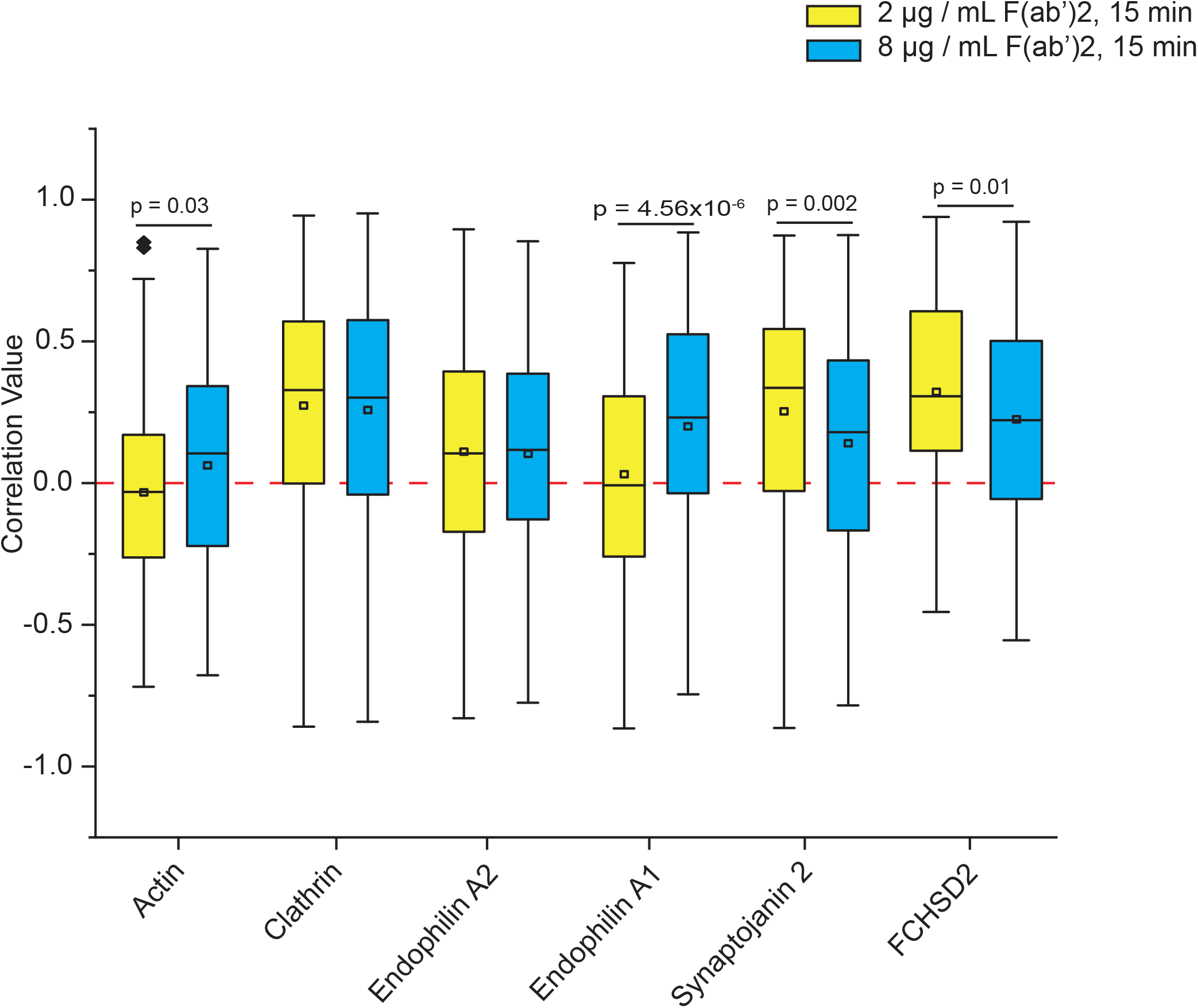
Analysis of coloclization of endocytic proteins and the BCR in cells stimulated with 2 or 8 μg/mL F(ab’)2. To identify endocytic proteins that are involved in generation of ccs on SRM structures we analyzed colocalization values of endocytic proteins (listed on x-axis) expressed with the BCR under conditions not expected to generate clathrin on SRM structures (2 μg/mL F(ab’)2, for 15 minutes) and conditions expected to generate mostly clathrin on SRM structures (8 μg/mL F(ab’)2, for 15 minutes). After a 15 minute stimulation, cells were fixed and imaged using TIRF microscopy. Small regions were extracted from each cell image and analyzed to calculate an average correlation value for each region as previously described in Larson BT et. al. The box plots represent all small regions analyzed for each cell and show the mean (square), median (line), 25/75 percentile range (box), and outliers with a coefficient value of 1.5. Actin was visualized using an Alexa Fluor 594 phalloidin stain. Clathrin, Endophilin A1 and A2 and Synaptojanin were all over-expressed with an mCherry fusion tag. And FCHSD2 was over-expressed with a GFP fusion tag. For co-localization with actin, clathrin, endophilin A1 and A2, and Synaptojanin, the BCR was over-expressed as a fusion protein with GFP (IgM-GFP). For colocalization analysis with FCHSD2-GFP, the BCR was stained using an Alexa Fluor 594 labeled F(ab) fragment.

**Supplemental Figure 5.**
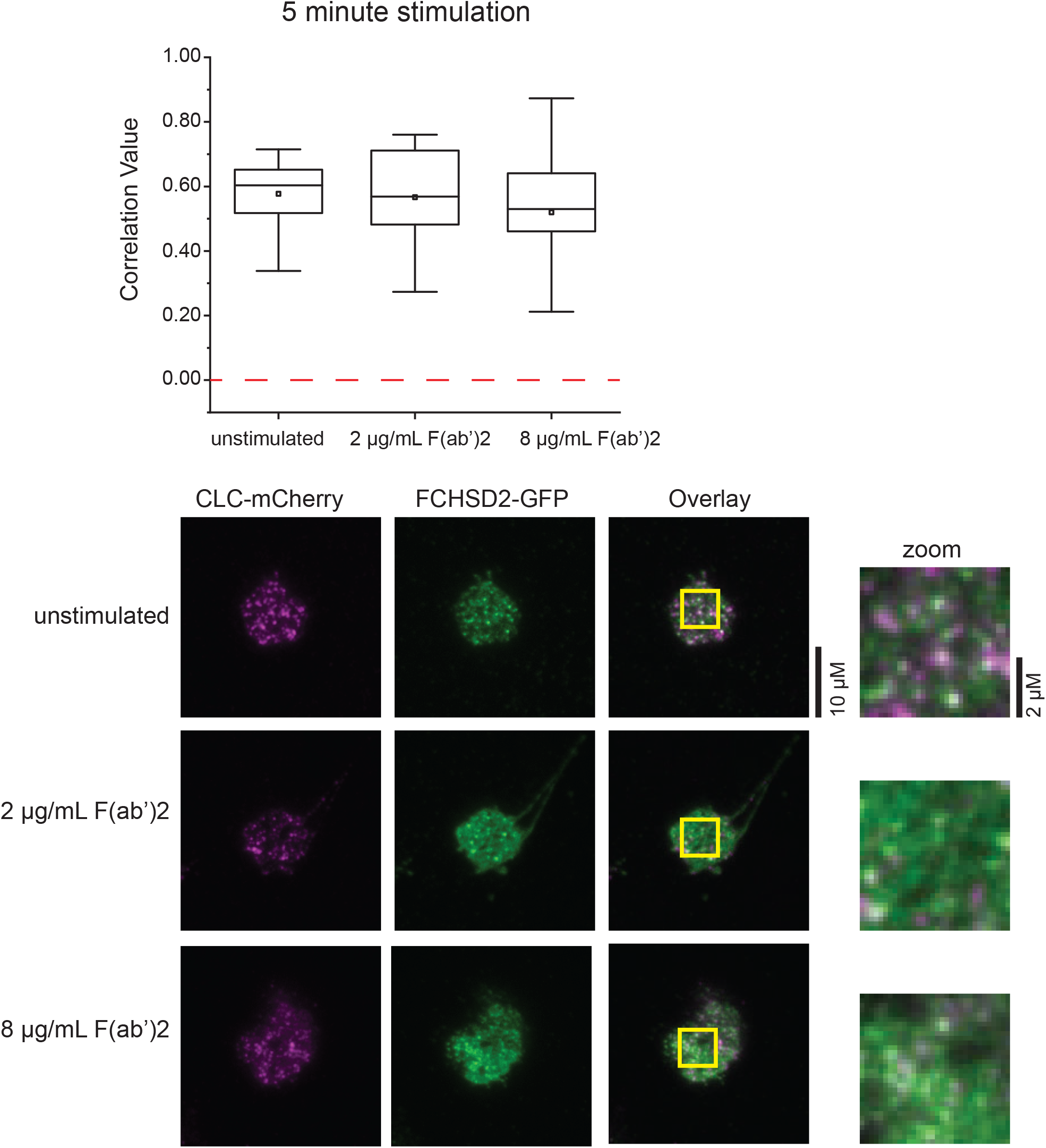
Colocalization analysis of clathrin and FCHSD2 in DG-75 cells. Colocalization analysis of FCHSD2 (FCHSD2-GFP overexpression) and clathrin (Clathrin Light Chain fused to mCherry overexpression) in DG-75 cells left unstimulated or stimulated with 2 or 8μg/mL F(ab’)2 for 5 minutes. The images below the graph show representative cells from each condition. Clathrin light chain is magenta and FHCSD2 is green.

**Supplemental Figure 6.**
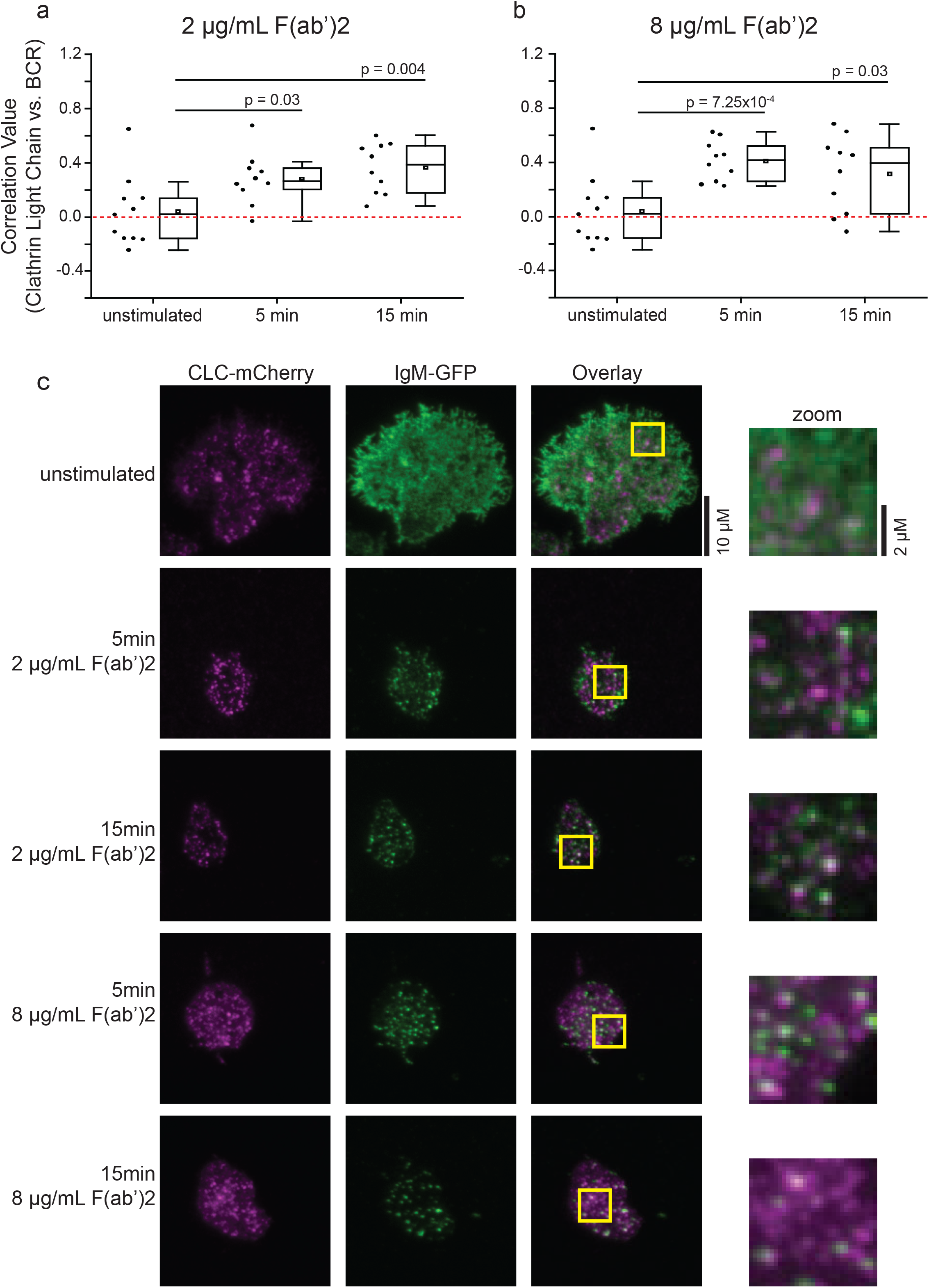
Colocalization analysis of clathrin and the BCR in DG-75 cells. (a) colocalization analysis of the BCR (IgM-GFP overexpression) and clathrin (clathrin light chain-mChery overexpression) in DG-75 cells stimulated with 2 μg/mL F(ab’)2 for 5 or 15 minutes. b) the same colocalization analysis done in part a is shown for cells treated with 8 μg/mL F(ab’)2 (c) representative TIRF microscopy images from the colocalization analysis presented in parts a and b. The zoomed images are of from the regions highlighted in yellow boxes. The box plots from parts a and b show the mean (square), median (line), 25/75 percentile range (box), and outliers with a coefficient value of 1.5 and data points (circles) from each cell.

**Supplemental Figure 7.**
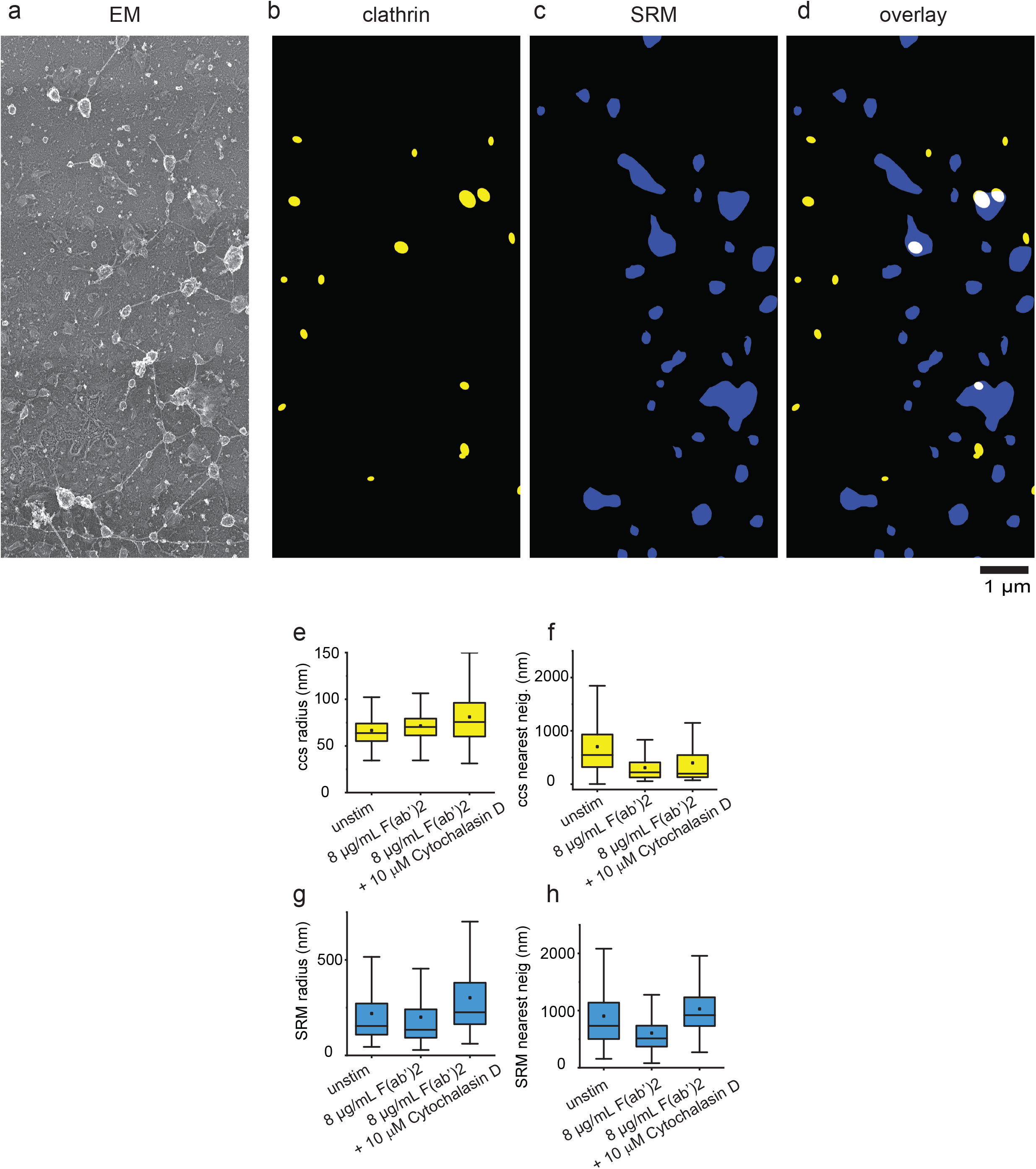
CCS and SRM radius and nearest neighbor distances following Cytochalasin D treatment. An example of an EM image (a) and corresponding segmented masks (b-d) that were used to measure changes in CCS radius (a), and (b) the nearest-neighbor distances between CCS (n = Unstim. 16 cells, 820 ccs structures, 15 min 8 μg/ml 12 cells, 2063 ccs structures, 10 μM Cytochalasin D 15 min 8 μg/ml 5 cells, 601 ccs structures). The same masks were also used to measure SRM radius (c), and nearest-neighbor distance (d) (n = Unstim. 16 cells, 708 SRM structures, 15 min 8 μg/ml 12 cells, 870 SRM structures, 10 μM Cytochalasin D 15 min 8 μg/ml 5 cells, 322 SRM structures).

